# The nutrient-sensing GCN2 signaling pathway is essential for circadian clock function by regulating histone acetylation under amino acid starvation

**DOI:** 10.1101/2022.12.08.519602

**Authors:** Xiaolan Liu, Yulin Yang, Yue Hu, Jingjing Wu, Chuqiao Han, Xihui Gan, Qiaojia Lu, Shaohua Qi, Jinhu Guo, Qun He, Yi Liu, Xiao Liu

**Author notes:** Corresponding author: Xiao Liu, Institute of Microbiology, Chinese Academy of Sciences, No.1 Beichen West Road, Chaoyang District, Beijing, 100101, China. equal contribution (Xiaolan Liu, Yulin Yang).

## Abstract

Circadian clocks are evolved to adapt to the daily environment changes under different conditions. The ability to maintain circadian clock functions in response to various stress and perturbations is important for organismal fitness. Here, we show that the nutrient sensing GCN2 signaling pathway is required for robust circadian clock function under amino acid starvation in *Neurospora*. The deletion of GCN2 pathway components disrupts rhythmic transcription of clock gene *frq* by suppressing WC complex binding at the *frq* promoter due to its reduced histone H3 acetylation levels. Under amino acid starvation, the activation of GCN2 kinase and its downstream transcription factor CPC-1 establish a proper chromatin state at the *frq* promoter by recruiting the histone acetyltransferase GCN-5. The arrhythmic phenotype of the GCN2 kinase mutants under amino acid starvation can be rescued by inhibiting histone deacetylation. Finally, genome-wide transcriptional analysis indicates that the GCN2 signaling pathway maintains robust rhythmic expression of metabolic genes under amino acid starvation. Together, these results uncover an essential role of GCN2 signaling pathway in maintaining robust circadian clock function in response to amino acid starvation and the importance of histone acetylation at the *frq* locus in rhythmic gene expression.

## Introduction

Circadian clocks are evolved to adapt to the daily environmental changes caused by the earth rotation (Bell-Pedersen et al., 2005; Dunlap & Loros, 2017; Johnson, Zhao, Xu, & Mori, 2017; Takahashi, 2017). Rhythmic gene expression allows different organisms to regulate their daily molecular, cellular, behavioral and physiological activities. The ability to maintain circadian clock function in response to various stress and perturbations is an important property of living systems (Bass, 2012; Hogenesch & Ueda, 2011). Although gene expression is sensitive to temperature changes, temperature compensation is a key feature of circadian clocks that maintain circadian period length at different physiological temperatures (Hu et al., 2021; Narasimamurthy & Virshup, 2021; Ode & Ueda, 2018). DNA damage and translation stress are known to reset circadian clock through the checkpoint kinase 2 signaling pathway in *Neurospora* and the ATM signaling pathway in mammalian cells (Diernfellner, Lauinger, Shostak, & Brunner, 2019; Oklejewicz et al., 2008; Pregueiro, Liu, Baker, Dunlap, & Loros, 2006). Cellular redox balance, including oxidative stress, regulates circadian clock by modulating CLOCK and NPAS2 activity (Nakahata et al., 2008; Rutter, Reick, Wu, & McKnight, 2001). Nutritional stress, such as high-fat diet, can disrupt the oscillating metabolites and behavioral circadian rhythms in mice (Eckel-Mahan et al., 2013; Kohsaka et al., 2007; Panda, 2016).

Starvation for all or certain amino acids leads to induced transcription followed by de-repression of genes in many amino acid biosynthetic pathways, referred as general amino acid control (GAAC) in yeast and cross-pathway control (CPC) in *Neurospora* (Hinnebusch, 2005). General control nonderepressible 2 (GCN2) kinase, called CPC-3 in *Neurospora*, is a serine/threonine kinase that functions as an amino acid sensor (Battu, Minhas, Mishra, & Khan, 2017; Efeyan, Comb, & Sabatini, 2015; Sattlegger, Hinnebusch, & Barthelmess, 1998). GCN2 is activated by accumulated uncharged tRNAs when intracellular amino acids are limited (Ramirez et al., 1992; Wek, Zhu, & Wek, 1995). Activated GCN2 phosphorylates the α subunit of eukaryotic initiation factor 2 (eIF2α) to translationally repress protein synthesis (Lyu, Yang, Zhao, & Liu, 2021; Sonenberg & Hinnebusch, 2009). Meanwhile, it also up-regulates the transcription activator GCN4, named CPC-1 in *Neurospora*, which activates amino acid biosynthetic and transport pathways (Ebbole, Paluh, Plamann, Sachs, & Yanofsky, 1991; Hinnebusch, 2005). Recently, it was shown that circadian clock control of GCN2 mediated eIF2α phosphorylation is necessary for rhythmic translation initiation in *Neurospora* (Ding, Lamb, Boukhris, Porter, & Bell-Pedersen, 2021; Karki et al., 2020). Although amino acid starvation is known to activate GCN2-GCN4 signaling pathway, how nutrient limitation, especially amino acid starvation, affects circadian clock function is not known.

Despite evolutionary distance in eukaryotes, circadian rhythms are controlled by transcription/translation-based negative feedback loops (Bell-Pedersen et al., 2005). *Neurospora crassa* was established as one of the best studied model systems for the molecular mechanism of eukaryotic circadian clocks (Cha, Zhou, & Liu, 2015; Dunlap & Loros, 2017; Heintzen & Liu, 2007). In the core *Neurospora* circadian negative feedback loop, two PAS-domain-containing transcription factors, White Collar-1 (WC-1) and WC-2 form a complex (WCC) and bind to the C-box of the *frq* promoter to activate its transcription (Cheng, Yang, & Liu, 2001; Crosthwaite, Dunlap, & Loros, 1997; Froehlich, Liu, Loros, & Dunlap, 2002). FRQ protein is translated from *frq* mRNA in the cytosol and progressively phosphorylated at about 100 phosphorylation sites, which plays a major role in determining circadian periodicity by regulating the FRQ-CK1 interaction (Baker, Kettenbach, Loros, Gerber, & Dunlap, 2009; Larrondo, Olivares-Yanez, Baker, Loros, & Dunlap, 2015; X. Liu et al., 2019; Y. Liu, Loros, & Dunlap, 2000; Tang et al., 2009). To close the negative feedback loop, FRQ forms a complex with its partner FRQ-interacting RNA helicase (FRH) to inhibit the activity of the WCC by promoting WC phosphorylation mediated by CK1 and CK2 (Q. He et al., 2006; Q. He & Y. Liu, 2005; Schafmeier et al., 2005; Wang, Kettenbach, Zhou, Loros, & Dunlap, 2019).

Chromatin structure and histone modification changes play important roles in regulating the transcription of circadian clock genes (Papazyan, Zhang, & Lazar, 2016; Takahashi, 2017; Zhu & Belden, 2020). In mammals, rhythmic H3K4me3, H3K9ac and H3K27ac modifications have been shown to be enriched at the promoter of clock genes and positively correlated with gene expression (Etchegaray, Lee, Wade, & Reppert, 2003; Katada & Sassone-Corsi, 2010; Koike et al., 2012). In *Neurospora*, rhythmic binding of WCC to the *frq* promoter is regulated by ATP-dependent chromatin remodeling factors, such as SWI/SNF complex, INO80 complex, the chromodomain helicase DNA-binding-1 (CHD1) and Clock ATPase (CATP) (Belden, Lewis, Selker, Loros, & Dunlap, 2011; Cha, Zhou, & Liu, 2013; Gai et al., 2017; Wang, Kettenbach, Gerber, Loros, & Dunlap, 2014). Histone chaperone FACT complex and histone modification enzymes, SET1, SET2 and RPD3 complex, have also been shown to affect *frq* transcription by regulating rhythmic histone compositions and modifications at the *frq* locus (X. Liu et al., 2017; Raduwan, Isola, & Belden, 2013; Sun et al., 2016). However, it is still unknown how chromatin structure is organized to allow rhythmic clock gene expression under nutrient limitation conditions.

In this study, we discovered that the disruption of the GCN2 (CPC-3) signaling pathway abolished robust circadian rhythms under amino acid starvation, which is important for rhythmic expression of metabolic genes. In the GCN2 signaling pathway mutants, amino acid starvation abolishes the rhythmic binding of WC-2 at the *frq* promoter by decreasing its histone H3 acetylation levels. Amino acid starvation activated CPC-3 and CPC-1, which re-established a proper chromatin state at the *frq* promoter by recruiting the histone acetyltransferase GCN-5 to allow rhythmic *frq* expression. Furthermore, the inhibition of histone deacetylases or disruption of the histone deacetylase can rescue the impaired circadian rhythm phenotypes under amino acid starvation, demonstrating the importance of rhythmic histone acetylation at the *frq* gene promoter for maintaining robust circadian gene expression.

## Results

### CPC-3 and CPC-1 are required for robust circadian rhythms under amino acid starvation

To investigate whether the nutrient sensing GCN2 pathway is involved in regulating circadian clock function under amino acid starvation, we created *cpc-3* and *cpc-1* knockout mutants. *cpc-3* and *cpc-1* encode for the *Neurospora* GCN2 and GCN4 homolog, respectively. As expected, the eIF2α phosphorylation and its induction by 3-aminotriazole (3-AT) treatment were completely abolished in the *cpc-3^KO^* strain (Figure 1–figure supplement 1A). 3-AT is an inhibitor of the histidine synthesis enzyme encoded by *his-3* which triggers amino acid starvation response (Natarajan et al., 2001). On the other hand, CPC-1 expression and its induction by 3-AT were eliminated in the *cpc-1 ^KO^* mutant (Figure 1–figure supplement 1B). The circadian conidiation rhythms of the *cpc-3^KO^* and *cpc-1 ^KO^* mutants were examined by race tube assays. Under a normal growth condition, *cpc-3* deletion had no effect on circadian period but *cpc-1* deletion led to period lengthening of ∼1.7 hr (Figures 1A). The long period of *cpc-1 ^KO^* strain could be rescued by the expression of Myc.CPC-1 (Figure 1–figure supplement 1C).

**Figure 1.**
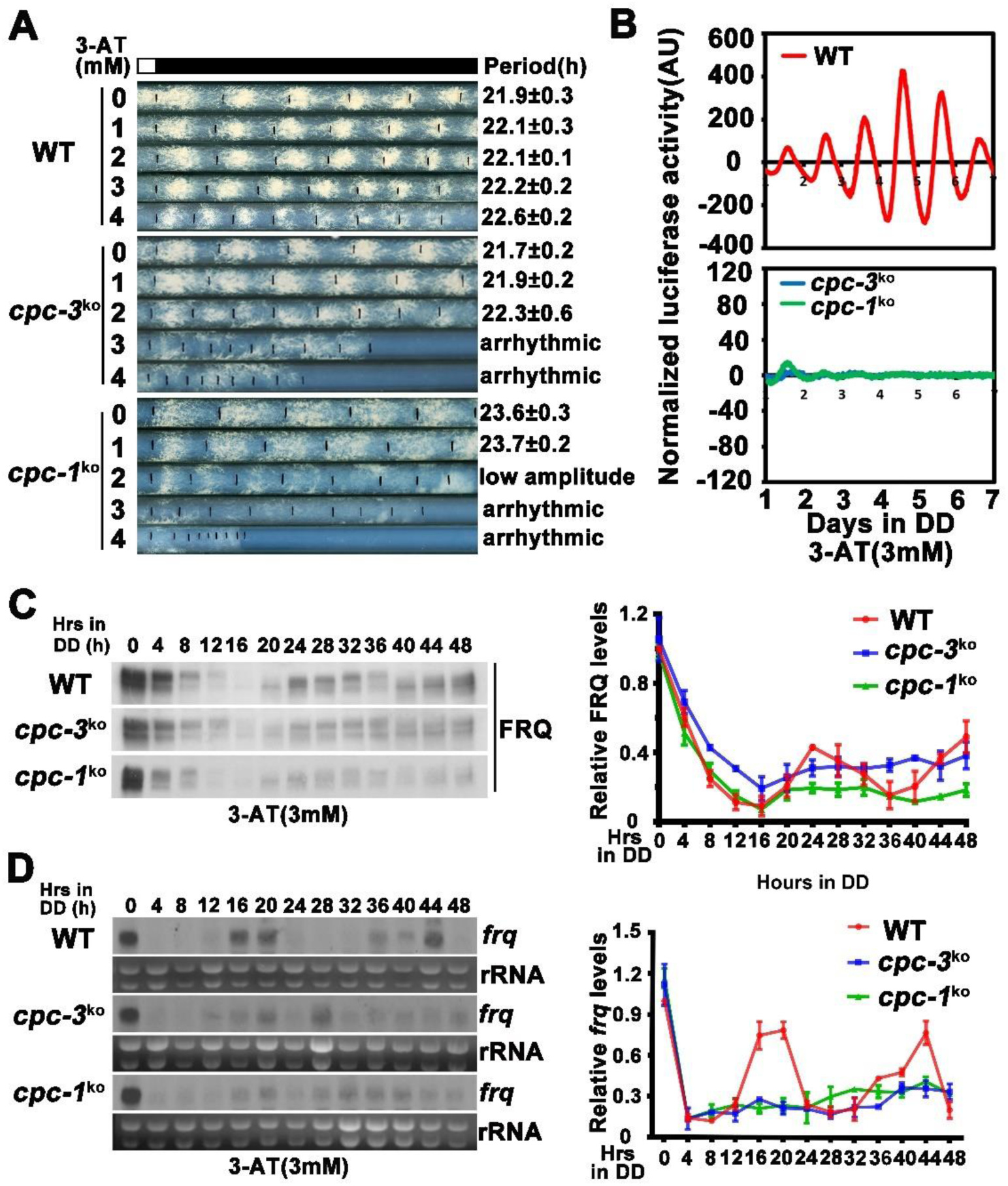
CPC-3 and CPC-1 are required for circadian rhythm by regulating rhythmic *frq* transcription in response to amino acid starvation. (A) Race tube assay showing that amino acid starvation (3-AT treatment) disrupted circadian conidiation rhythm of the *cpc-3^KO^* and *cpc-1^KO^* strains. (B) Luciferase reporter assay showing that amino acid starvation disrupted rhythmic expression of *frq* promoter driven luciferase of the *cpc-3^KO^* and *cpc-1^KO^* strains. A *frq-luc* transcriptional fusion construct was expressed in *cpc-3^KO^* and *cpc-1^KO^* strains grown on the FGS-Vogel’s medium with the indicated concentrations of 3-AT, and the luciferase signal was recorded using a LumiCycle in constant darkness (DD) for more than 7 days. Normalized data with the baseline luciferase signals subtracted are shown. (C) Western blot showing that amino acid starvation dampened rhythmic expression of FRQ protein of the *cpc-3^KO^* and *cpc-1^KO^* strains at the indicated time points in DD. The left panel showing that protein extracts were isolated from WT, *cpc-3^KO^* and *cpc-1^KO^* strains grown in a circadian time course in DD and probed with FRQ antibody. The right panel showing that the densitometric analyses of the results of three independent experiments. (D) Northern blot showing that amino acid starvation dampened rhythmic expression of *frq* mRNA of the *cpc-3^KO^* and *cpc-1^KO^* strains at the indicated time points in DD. The densitometric analyses of the results from three independent experiments are shown on the right panel.

3-AT is an inhibitor of the histidine synthesis enzyme encoded by *his-3* which triggers amino acid starvation response (Natarajan et al., 2001). To investigate how amino acid starvation affects circadian clock function, we treated the wild-type, *cpc-3 ^KO^* and *cpc-1 ^KO^* strains with different concentrations of 3-AT. As shown in Figure 1A, although 3 and 4 mM 3-AT resulted in modest inhibition of the WT growth rate, robust circadian conidiation rhythms were maintained. For the *cpc-3 ^KO^* and *cpc-1 ^KO^* strains, however, 3 and 4 mM 3-AT resulted in severe inhibition of growth rates and the loss of circadian conidiation rhythms. To exclude the effect of 3-AT on other target genes, we examined the circadian rhythm of the *his-3^-^* strain, which contains a single mutation in a gene required for histidine synthesis and could not grow in the medium without histidine. Race tube assay showed that the *his-3^-^* strain grew normally and exhibited a robust circadian conidiation rhythm in the presence of histidine (1.0*10^-2^ mg/mL). Although the same amount of histidine addition could rescue the growth of the *cpc-3 ^KO^ his-3^-^* strain, it could not rescue its circadian conidiation rhythm (Figure 1–figure supplement 2A), indicating that CPC-3 is required for robust circadian rhythms under histidine starvation stress.

To confirm the loss of circadian rhythm at the molecular level, we introduced a *frq* promoter driven luciferase reporter into the *cpc-3^KO^* and *cpc-1 ^KO^* strains. As shown in Figures 1B and Figure 1–figure supplement 3A, the robust circadian rhythm of luciferase activity seen in the wild-type strains were severely dampened to arrhythmic in the *cpc-3^KO^* and *cpc-1 ^KO^* strains at 3 mM 3-AT treatment. Consistent with these results, western blot analysis showed that, after the initial light/dark transition, no obvious rhythms of FRQ abundance and its phosphorylation profile were seen in the *cpc-3^KO^* and *cpc-1^KO^* strains in the presence of 3-AT (Figures 1C and Figure 1–figure supplement 3B). Northern blot analysis showed that the circadian rhythms of *frq* mRNA in the *cpc-3^KO^* and *cpc-1 ^KO^* strains were also abolished in the presence of 3-AT and the levels of *frq* mRNA were constantly low in DD (Figures 1D and Figure 1–figure supplement 3C). Together, these results suggest that the GCN2 pathway is required for a functional clock by regulating rhythmic *frq* transcription in response to amino acid starvation.

### CPC-3 and CPC-1 are required for rhythmic WCC binding in response to amino acid starvation

Since 3-AT treatment resulted in low *frq* mRNA levels in the *cpc-3^KO^* and *cpc-1^KO^* strains (Figure 1D), we first examined the expression levels of WC-1 and WC-2 and found that their levels were higher in the mutants than the wild-type strain at different time points (Figures 2A and 2B). We then performed WC-2 ChIP assays to examine whether WC binding to the *frq* promoter was affected. As shown in Figure 2–figure supplement 1A, WC-2 rhythmically bound to *frq* C-box in the WT, *cpc-3^KO^* and *cpc-1 ^KO^* strains without 3-AT treatment. However, 3-AT treatment resulted in constant low levels of WC-2 binding to the *frq* C-box during a circadian cycle in both *cpc-3^KO^* and *cpc-1 ^KO^* strains (Figures 2C and 2D). These results indicate that the loss of circadian rhythms in the *cpc-3^KO^* and *cpc-1 ^KO^* strains under amino acid starvation is caused by loss of rhythmic *frq* transcription due to impaired WC binding at the *frq* promoter.

**Figure 2.**
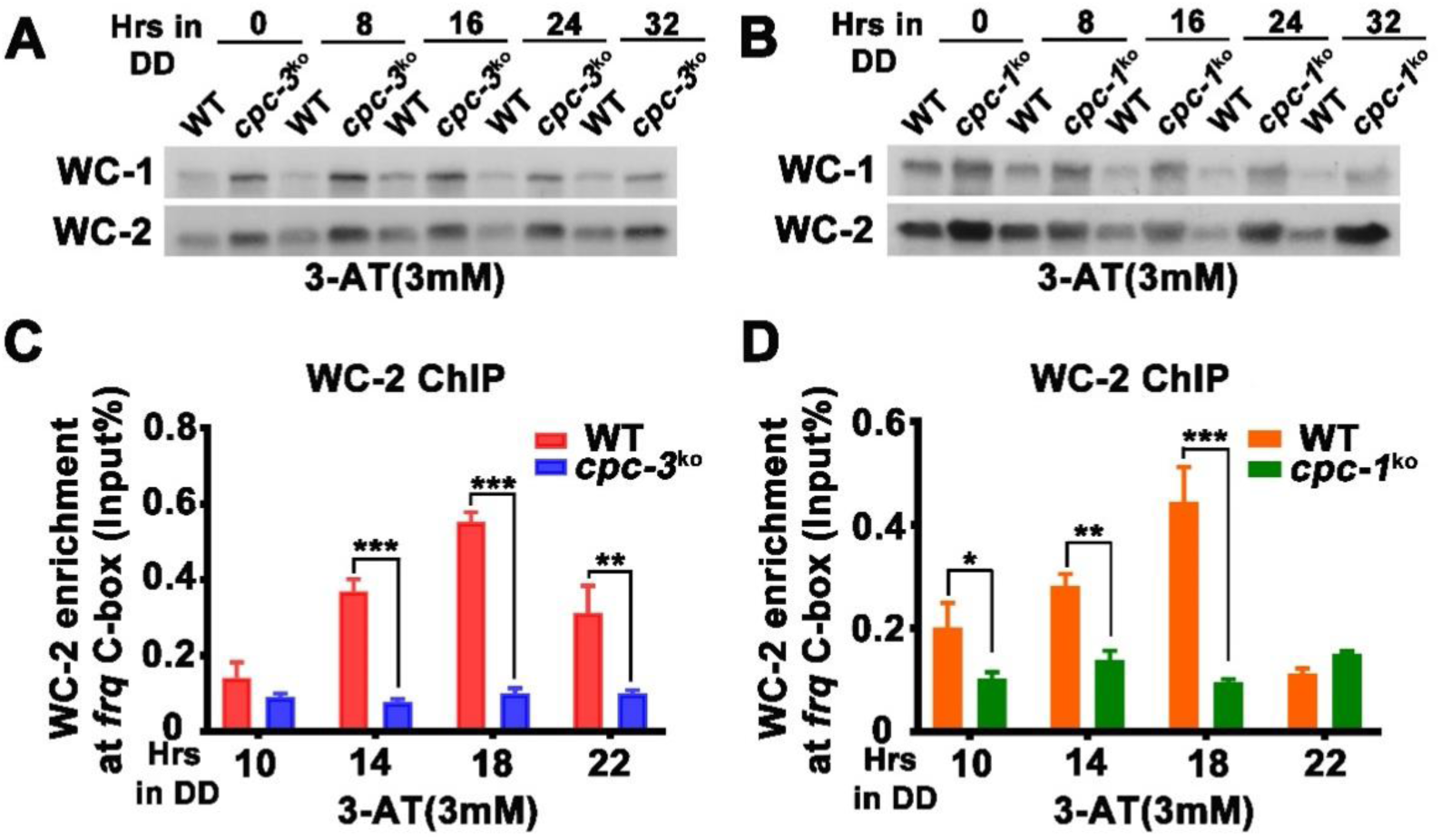
CPC-3 and CPC-1 are required for rhythmic WCC binding in response to amino acid starvation. (A-B) Western blot assay showing that WCC protein levels were elevated in the *cpc-3^KO^* (A) and *cpc-1^KO^* (B) strains after 3 mM 3-AT treatment. Protein extracts were isolated from WT, *cpc-3^KO^* and *cpc-1^KO^* strains grown in the indicated time points in DD and probed with WC-1 and WC-2 antibodies. (C-D) ChIP assay showing that amino acid starvation disrupted rhythmic WC-2 binding at the promoter of *frq* gene in the *cpc-3^KO^* (C) or *cpc-1^KO^* strains(D). Samples were grown for the indicated number of hours in DD prior to harvesting and processing for ChIP using WC-2 antibody. Occupancies were normalized by the ratio of ChIP to Input DNA. Error bars indicate standard deviation (n = 3). *p < 0.05; **p < 0.01; ***P<0.001; Student’s t test was used.

Because WC phosphorylation inhibits the transcriptional activation activity of the WC complex (Q. He et al., 2006; Q. Y. He & Y. Liu, 2005; Lee, Loros, & Dunlap, 2000; Schafmeier et al., 2005), we also examined WC phosphorylation profiles and found that 3-AT treatment resulted in hypophosphorylation of WC-1 and WC-2 (which is normally associated with WC activation) in the *cpc-3^KO^* and *cpc-1 ^KO^* strains (Figure 2–figure supplement 1B and 1C). Thus, their reduced WC binding at the *frq* promoter is not caused by WC hyper-phosphorylation.

### CPC-1 is required for maintenance of chromatin structure in response to amino acid starvation

The low WC-2 binding at the *frq* promoter promoted us to examine the chromatin structure of the *frq* promoter. We first performed ChIP assay to examine the histone and its acetylation levels at the *frq* promoter in the WT strain. Although amino acid starvation had little effect on the histone H2B levels (Figure 3A), it significantly decreased histone H3 acetylation levels (H3 acetylated at the N-terminus) at the *frq* promoter at high concentrations of 3-AT (Figure 3B). These results suggest that amino acid starvation can affect *frq* transcription by reducing the histone acetylation levels at the *frq* promoter.

**Figure 3.**
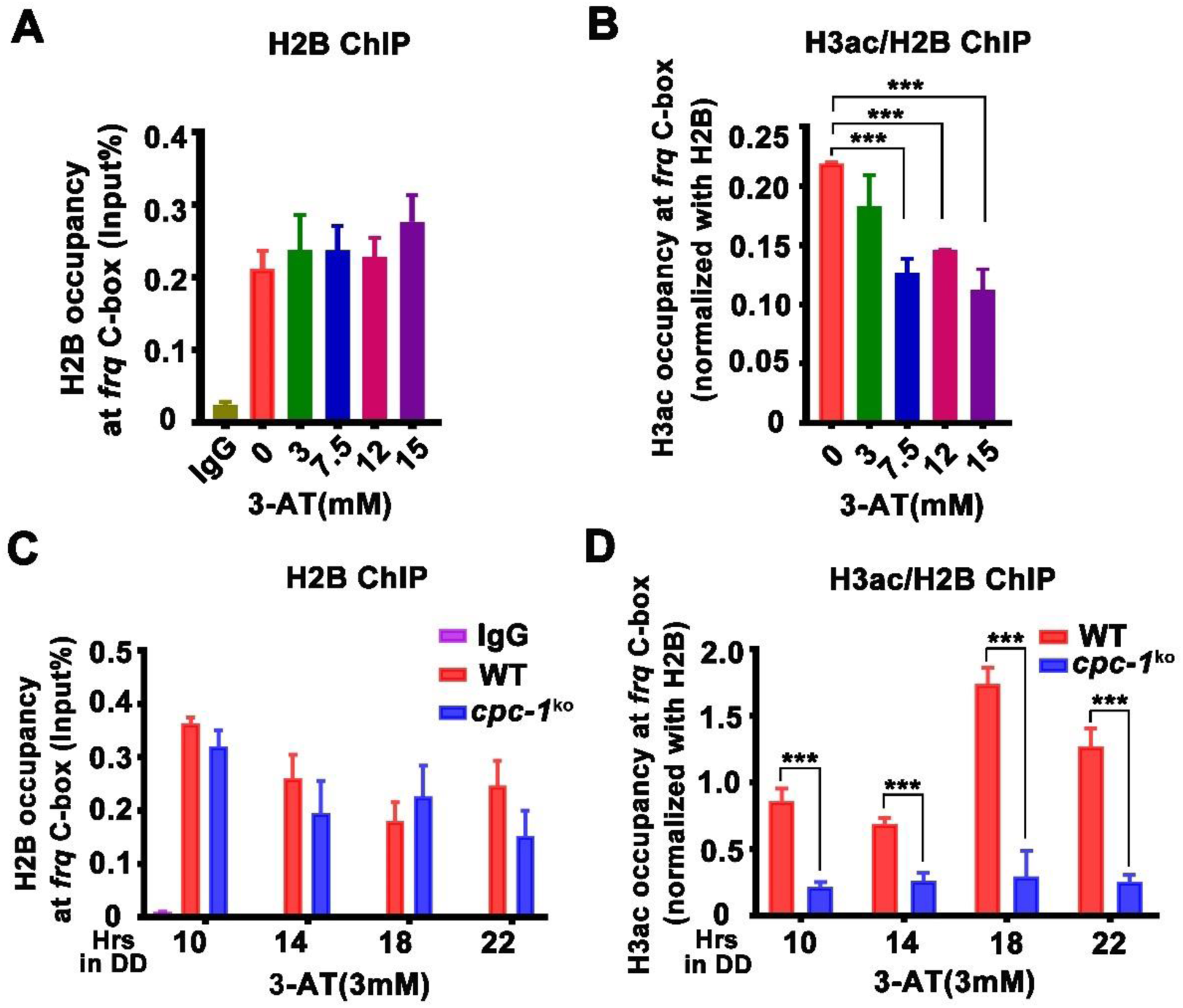
CPC-1 is required for maintenance of chromatin structure in response to amino acid starvation. (A-B) ChIP assay showing that amino acid starvation slightly increased histone H2B levels (A) and dramatically decreased histone H3ac levels (B) at the promoter of *frq* gene in the WT strain at DD18 at the indicated concentration of 3-AT. Relative H3ac levels were normalized with H2B levels. (C-D) ChIP assay showing that amino acid starvation slightly increased histone H2B levels (C) and dramatically decreased histone H3ac levels (D) at the promoter of *frq* gene in the *cpc-1^KO^* strain at the indicated time points in DD. Error bars indicate standard deviations (n=3). ***P<0.001; Student’s t test was used.

We then examined whether CPC-3 and CPC-1 signaling pathway was involved in regulating chromatin structure at the *frq* promoter in response to amino acid starvation. Histone H2B and H3ac ChIP assays at different circadian time points showed that histone H3ac levels were slightly decreased in the *cpc-1^KO^* strain compared to the WT strain under normal condition (Figure 2–figure supplement 1D). H2B levels were not markedly different between the WT and *cpc-1^KO^* strains in the presence of 3 mM 3-AT (Figure 3C). However, the relative histone H3ac levels are very different in these two strains in the presence of 3 mM 3-AT: it was rhythmic with a peak at DD18 in the WT strain but was dramatically low in the *cpc-1^KO^* strain at different time points in DD (Figure 3D). These results indicate that CPC-1 is required for maintaining the proper histone acetylation status at the *frq* promoter under amino acid starvation. The low H3ac level at the *frq* promoter, which is critical for transcription activation, results in constant low WCC binding and arrhythmic *frq* transcription in the *cpc-1^KO^* strain.

### CPC-1 recruits GCN-5 to activate *frq* transcription in response to amino acid starvation

To determine how CPC-1 is involved in regulating histone acetylation levels at the *frq* locus, we expressed Myc-tagged CPC-1 in the WT strain and examined the occupancy of Myc.CPC-1 at the *frq* promoter by ChIP assays. As shown in Figure 4A, Myc.CPC-1 was found to be rhythmically enriched at the *frq* promoter in DD, peaking at ∼DD14, a time point that is near the *frq* mRNA peak. Co-immunoprecipitation (co-IP) assay showed that CPC-1 did not associate with WC-1 or WC-2 (Figure 4–figure supplement 1A), suggesting that CPC-1 and WCC bind independently to the *frq* promoter.

**Figure 4.**
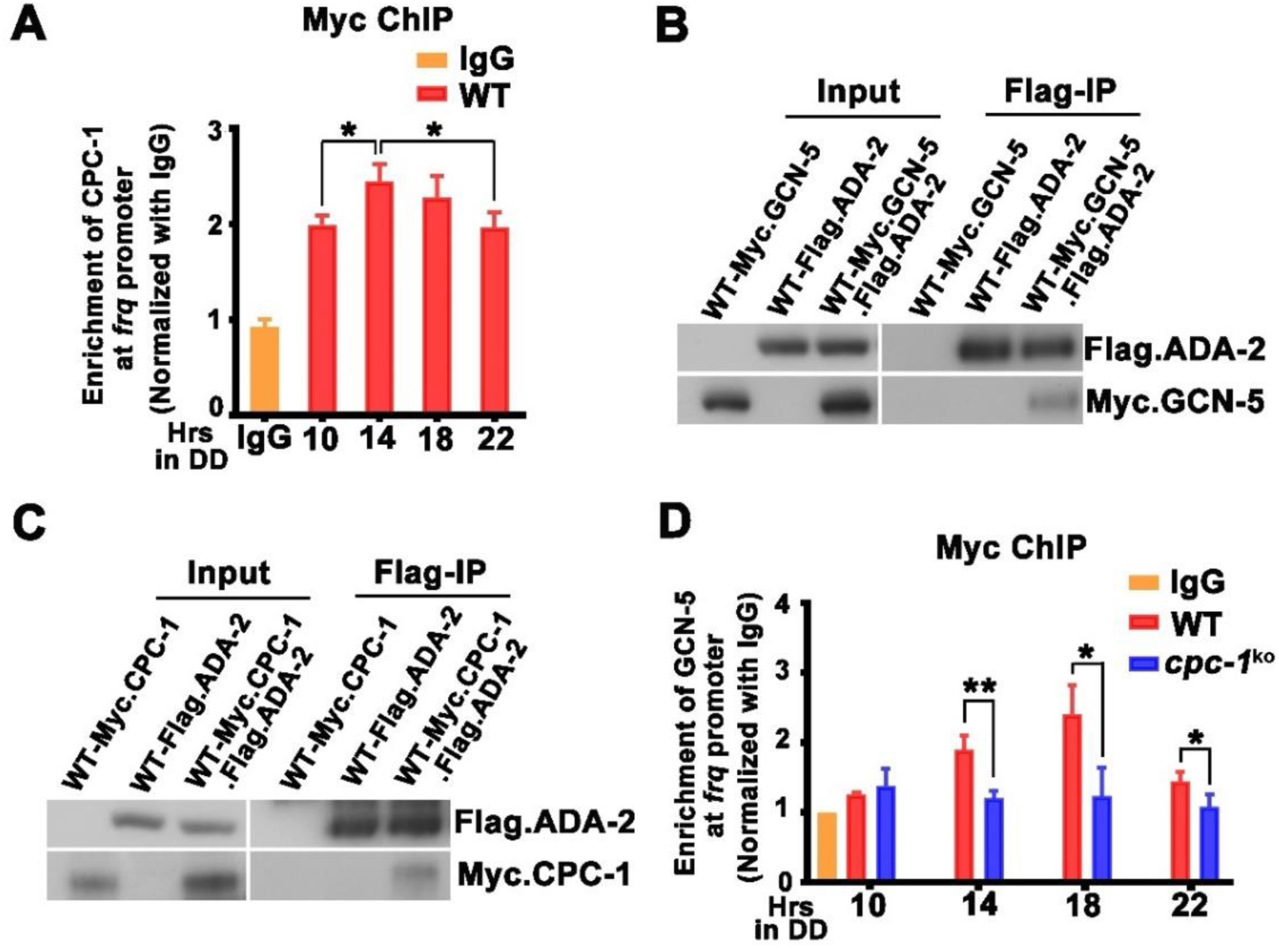
CPC-1 recruits GCN-5 to activate *frq* transcription in response to amino acid starvation. (A) ChIP assay showing that CPC-1 rhythmically bound at the promoter of *frq* gene. Myc.CPC-1 was expressed in the WT strain grown for the indicated number of hours in DD. Samples were crosslinked with formaldehyde and harvested for ChIP using Myc antibody. Myc ChIP occupancies were normalized with IgG. (B) IP assay showing that Flag.ADA-2 interacted with Myc.GCN-5. Flag.ADA-2 and Myc.GCN-5 were co-expressed in the WT strain and immunoprecipitation was performed using Flag antibody. (C) IP assay showing that Flag.ADA-2 interacted with Myc.CPC-1. Flag.ADA-2 and Myc.CPC-1 were co-expressed in the WT strain and immunoprecipitation was performed using Flag antibody. (D) ChIP assay showing that rhythmic GCN-5 binding at the promoter of *frq* gene was dampened in the *cpc-1^KO^* strain. Samples were grown for the indicated number of hours in DD prior to harvesting and processing for ChIP as described in (A). Error bars indicate standard deviations (n=3). *p < 0.05; **p < 0.01; Student’s t test was used.

How does GCN2 signaling pathway regulate histone acetylation in response to amino acid starvation? The yeast GCN4 was previously shown to recruit histone acetyltransferase GCN5 containing (Spt-Ada-Gcn5 acetyltransferase) SAGA complex to selective gene promoters, likely through its physical interaction with the ADA2 subunit (Barlev et al., 1995; Belotserkovskaya et al., 2000; Drysdale et al., 1998; Kuo, vom Baur, Struhl, & Allis, 2000). To determine this possibility, we performed co-immunoprecipitation assay to check the interaction between CPC-1 and the SAGA complex in *Neurospora* in strains expressing the epitope-tagged *Neurospora* SAGA homologs. As shown in Figure 4B, Myc-tagged GCN-5 was efficiently immunoprecipitated by the Flag-tagged ADA-2, indicating the existence of a SAGA complex in *Neurospora*. Importantly, Myc.CPC-1 was also found to associate specifically with Flag.ADA-2 (Figure 4C), suggesting that CPC-1 can recruit the SAGA complex to the *frq* promoter through its ADA-2 subunit to regulate histone acetylation levels at the *frq* locus. Furthermore, immunoprecipitation assays showed that WC-1 and WC-2 were also interact with Myc.GCN-5 (Figure 4–figure supplement 1B). These results suggest that CPC-1 can regulate the histone acetylation by recruiting the SAGA complex to the *frq* promoter.

To confirm if CPC-1 can recruit GCN5 to the *frq* promoter, we performed ChIP assay to examine the occupancy of GCN-5 at the *frq* promoter. As shown in Figure 4D, Myc-tagged GCN-5 rhythmically bound at the *frq* promoter in DD in the wild-type strain but its binding was constantly low in the *cpc-1^KO^* strain, suggesting that CPC-1 rhythmically recruit GCN-5 containing SAGA complex to the *frq* promoter to allow rhythmic histone acetylation levels to allow for rhythmic WC-2 binding and thus rhythmic *frq* transcription.

### GCN-5 is required for rhythmic H3ac at the *frq* promoter

GCN-5 was previously shown to regulate light induction and oxidative stress response in *Neurospora* (Grimaldi et al., 2006; Qi et al., 2018), but its role in circadian clock is unclear. To determine the function of GCN-5 in the circadian clock, we created the *gcn-5^KO^* strain and found that it exhibited a slow growth and lacked a conidiation rhythm (Figure 5A). To determine its circadian clock phenotype at the molecular level, we introduced the FRQ-LUC reporter (luciferase fused at the C terminus of the FRQ protein) into the *gcn-5^KO^* strain (Larrondo et al., 2015). As shown in Figure 5B, a robust circadian rhythm of luciferase activity was seen in a wild-type strain but was quickly dampened after one day in DD and became arrhythmic in the *gcn-5^KO^* strain. Western blot analysis showed that, after the initial light/dark transition, the rhythmic FRQ abundance and phosphorylation were significantly dampened in the mutant (Figure 5C). Northern blot analysis showed that the circadian rhythms of *frq* mRNA were also severely dampened in the *gcn-5^KO^* strain (Figure 5D). Together, these results indicate that GCN-5 is critical for circadian clock function.

**Figure 5.**
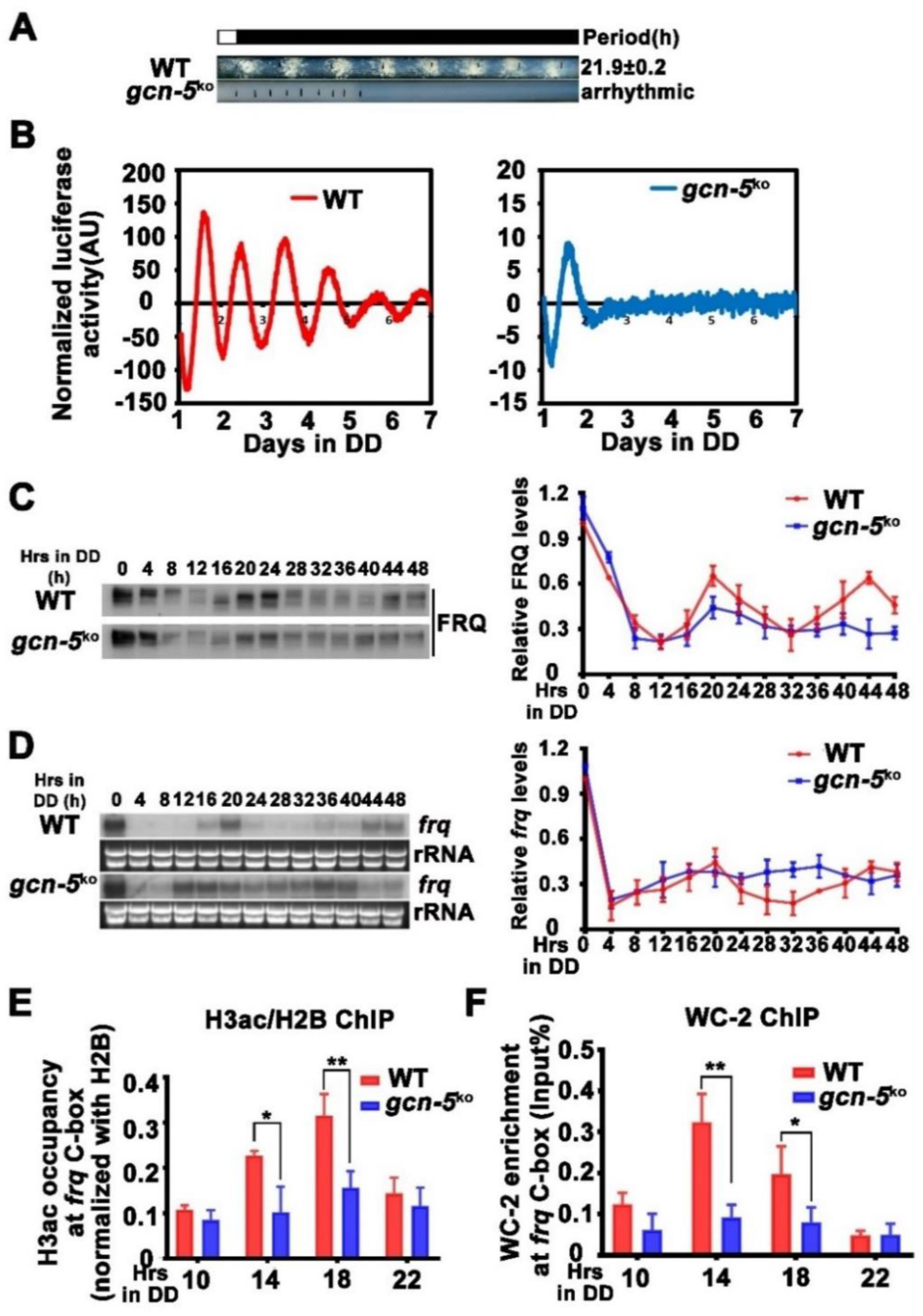
GCN-5 is required for rhythmic chromatin structure changes at the *frq* promoter. (A) Race tube assay showing that the conidiation rhythm in *gcn-5^KO^* strains was lost compared with wild-type strains. (B) Luciferase assay showing that the luciferase activity rhythm was impaired in the *gcn-5^KO^* strain after one day transition from light to dark. A FRQ-LUC translational fusion construct was expressed in WT and *gcn-5^KO^* strains, and the luciferase signal was recorded in DD for more than 7 days. Normalized data with the baseline luciferase signals subtracted are shown. (C) Western blot assay showing that rhythmic expression of FRQ protein was dampened in the *gcn-5^KO^* strain. (D) Northern blot assay showing that rhythmic expression of *frq* mRNA was dampened in the *gcn-5^KO^* strain. (E) ChIP assay showing decreased histone H3ac levels at the promoter of *frq* gene in the *gcn-5^KO^* strain at the indicated time points in DD. Relative H3ac levels were normalized with H2B levels. (F) ChIP assay showing decreased WC-2 levels at the promoter of *frq* gene in the *gcn-5^KO^* strain at the indicated time points in DD. Error bars indicate standard deviations (n=3). *p < 0.05; **p < 0.01; Student’s t test was used.

ChIP assays showed that the H3ac levels were significantly decreased at the *frq* promoter, and its rhythmic occupancy was severely dampened in the *gcn-5^KO^* strain compared with the WT strain (Figure 5E). Furthermore, the rhythmic WC-2 binding at the *frq* C-box of the WT strain was dramatically reduced in the *gcn-5^KO^* strain in DD (Figure 5F). These results indicate that GCN-5 is critical for circadian clock function by regulating rhythmic chromatin structure changes to allow rhythmic WC-2 binding at the *frq* promoter to drive rhythmic *frq* transcription.

Since ADA-2 interacts with GCN-5 and is a subunit of the SAGA complex, we also created *Neurospora ada-2^KO^* strains and examined the function of the ADA-2 in the circadian clock. As expected, we found that circadian phenotypes and *frq* expression of the *ada-2^KO^* strain were very similar to those of the *gcn-5^KO^* strain (Figure 5–figure supplement 1A-C). Together, these results demonstrate the importance of GCN-5 and ADA-2 in the *Neurospora* circadian clock function.

### Elevated histone acetylation partially rescues circadian clock defects caused by amino acid starvation

Our results above suggest that amino acid starvation results in low histone acetylation levels at the *frq* promoter, which impairs the WC-2 binding and rhythmic *frq* transcription in the *cpc-3 ^KO^* and *cpc-1^KO^* mutants. To further confirm this conclusion, we hypothesized that the circadian clock defects should be rescued by elevated histone acetylation levels. Trichostatin A (TSA) is a histone deacetylase (HDACs) inhibitor, which can inhibit HDACs activity and increase the histone acetylation levels in *Neurospora* (Selker, 1998). We treated the WT strain with different concentrations of 3-AT and TSA. We found that high concentrations of 3-AT lengthened the circadian period of the WT strain to ∼24 hours (Figure 6A), but high concentrations of TSA resulted in a slight period shortening to 21 hours (Figure 6B). When TSA was used together with a high concentration of 3-AT (7.5mM), the long period phenotype can be gradually rescued by increasing TSA concentration (Figure 6C), consistent with our conclusion that the histone acetylation level changes is responsible for the circadian clock defects caused by amino acid starvation.

**Figure 6.**
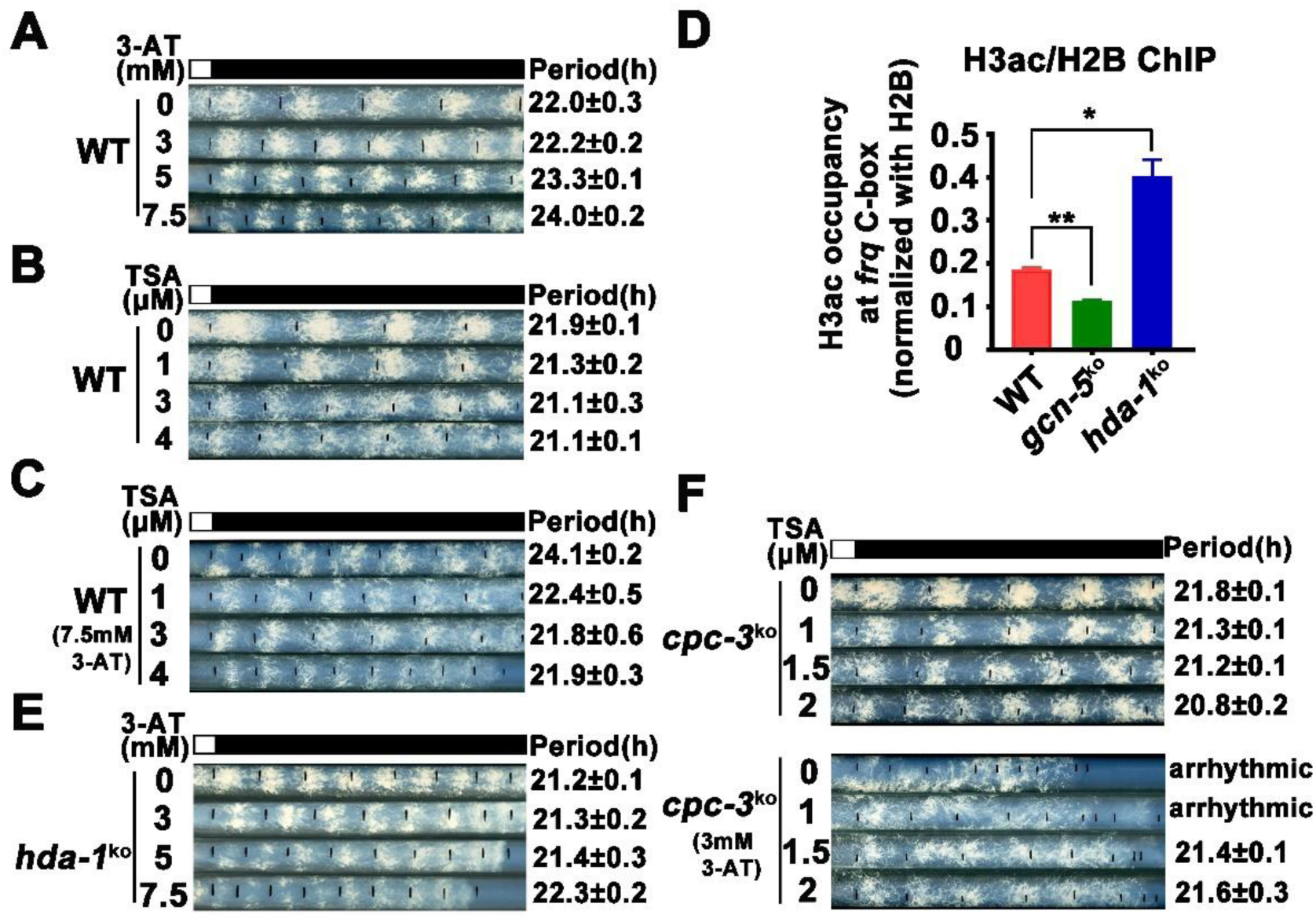
Elevated histone acetylation partially rescues impaired circadian rhythm caused by amino acid starvation stress. (A) Race tube assay showing that high concentrations of 3-AT treatment elongated circadian conidiation period of WT strain. (B) Race tube assay showing that high concentrations of TSA treatment shortened circadian conidiation period of WT strain. (C) TSA treatment rescued prolonged circadian period of WT strain caused by 3-AT treatment. WT strains were grown on the race tube medium containing 7.5mM 3-AT and different indicated concentrations of TSA in DD. (D) ChIP assay showing that H3ac levels were decreased in *gcn-5^KO^* strains and increased in *hda-1^KO^* strains at the promoter of *frq* gene. Error bars indicate standard deviations (n=3). *p < 0.05; **p < 0.01; Student’s t test was used. (E) The *hda-1^KO^* strain exhibited near normal circadian period in the presence of 3-AT. *hda-1^KO^* strains were grown on the race tube medium containing the indicated concentrations of 3-AT in DD. (F) TSA treatment rescued the impaired circadian rhythm of *cpc-3^KO^* strain caused by 3-AT treatment. *cpc-3^KO^* strains were grown on the race tube medium containing 3 mM 3-AT and different indicated concentrations of TSA in DD.

GCN-5 is the major histone acetyltransferase responsible for histone acetylation at *frq* locus in response to amino acid starvation. On the other hand, the histone deacetylase HDA-1 was previously reported as a major histone deacetylase that can antagonize and compete with GCN-5 for recruitment to promoters to deacetylate H3 (Islam, Turner, Menzel, Malo, & Harkness, 2011; Vogelauer, Wu, Suka, & Grunstein, 2000). ChIP assay showed that H3ac levels were indeed significantly decreased in the *gcn-5^KO^* strain and were markedly increased in the *hda-1^KO^* strains at the *frq* promoter region (Figure 6D), indicating that HDA-1 is responsible for histone deacetylation at the *frq* locus. Race tube assays showed that in contrast to period lengthening in the WT strain by 3-AT (Figure 6A), the *hda-1^KO^* strain exhibited near normal circadian period even in the presence of high 3-AT concentrations (Figure 6E), suggesting that elevated histone deacetylation can partially rescue the circadian clock defects in response to amino acid starvation. To further confirm our conclusion, we treated the *cpc-3^KO^* strains with different concentrations of TSA and found that the arrhythmic circadian conidiation rhythm caused by 3-AT treatment could be partially rescued by TSA treatments (Figure 6F). Together, these results strongly suggested that the amino acid starvation-induced clock defects in the GCN2 signaling pathway mutants are largely due to the decreased histone acetylation at the *frq* promoter, which prevents efficient WC-2 binding and rhythmic *frq* transcription.

### Rhythmic expression of CPC-1 activated metabolic genes under amino acid starvation

Circadian clock has been shown to control metabolic processes and rhythmic transcription of metabolic genes (Baek et al., 2019; Hurley et al., 2014). To determine the role of the GCN2 pathway in controlling gene expression under amino acid starvation, we performed RNA-seq experiments to analyze the genome-wide mRNA levels in the WT and *cpc-1^KO^* strains in the presence of 3-AT (12 mM). As shown in Figure 7A, 22.1% of genes were found to be up-regulated and 11.2% of genes were down-regulated in the WT strain after 3-AT treatment compared with normal condition. Specifically, genes involved in the regulation of oxidoreductase and amino acid metabolism were particularly enriched and were mostly up-regulated during the amino acid starvation (Figure 7B). However, the differential expression of the 148 up-regulated and 127 down-regulated genes found in the WT strain were abolished in the *cpc-1^KO^* strain after 3-AT treatment, suggesting that these genes were regulated by CPC-1 under amino acid starvation (Figures 7D and Figure 7–figure supplement 1A-B). Among them, genes involved in amino acid and vitamin metabolism were particularly enriched and were mostly up-regulated during the amino acid starvation (Figure 7E). For example, amino acid synthesis genes *his-3* (NCU03139), *ser-2* (NCU01439) and *trp-3* (NCU08409) were up-regulated in the WT strain, but unchanged in the *cpc-1^KO^* strain under amino acid starvation (Figures 7A and 7C). Importantly, these three genes were all exhibited rhythmic expression in DD in the WT but not in the *cpc-1^KO^* strain (Figure 7F), suggesting that CPC-1-activated metabolic genes under amino acid starvation were regulated by circadian clock. Together, these results suggest that the GCN2 signaling pathway maintained robust circadian clock function and rhythmic expression of metabolic genes under amino acid starvation.

**Figure 7.**
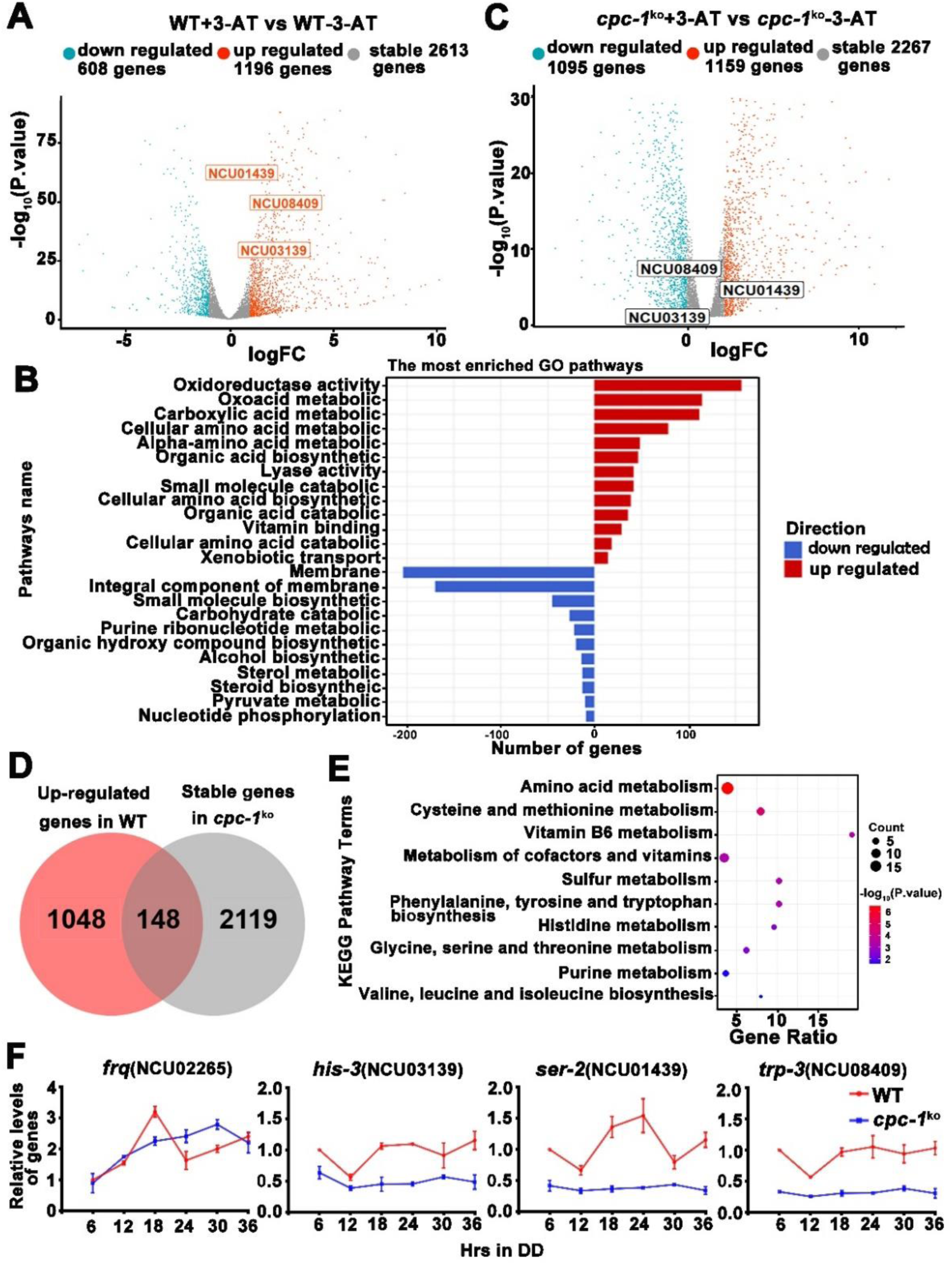
Circadian control of CPC-1-activated metabolic genes under amino acid starvation. (A) Comparison of the transcript expression profiles of the WT strains with and without 12 mM 3-AT treatment. (B) Gene functional enrichment analysis based on the mRNA level changes for the up- and down-regulated genes in the WT strains with and without 12 mM 3-AT treatment. (C) Comparison of the transcript expression profiles of the *cpc-1^KO^* strains with and without 12 mM 3-AT treatment. (D) Pie charts showing the overlaps of up-regulated genes in the WT strain, but stable genes in the *cpc-1^KO^* strains after 12 mM 3-AT treatment. (E) Gene functional enrichment analysis based on the mRNA level changes for the overlaps of up-regulated genes in the WT strain, but stable genes in the *cpc-1^KO^* strains after 12 mM 3-AT treatment. (F) RT-qPCR analysis showing that amino acid synthetic genes *his-3* (NCU03139), *ser-2* (NCU01439) and *trp-3* (NCU08409) were activated by CPC-1 and were rhythmic expressed. The primers used for RT-qPCR were shown in Figure 7–Table supplement 1.

## Discussion

Circadian clock drives robust rhythmic gene expression and activities at different environmental and nutrient conditions. Here, we discovered that the nutrient sensing GCN2 pathway plays an unexpected role in maintaining the *Neurospora* circadian clock in response to amino acid starvation stress. Amino acid starvation activates the CPC-3 and CPC-1 signaling pathway and CPC-1 specifically recruits the histone acetyltransferase GCN-5 containing SAGA complex to the *frq* promoter to increase the histone acetylation levels, which permits rhythmic WC-2 binding. Therefore, under amino acid starvation, the disruption of the CPC-3 and CPC-1 signaling pathway results in decreased histone acetylation levels, reduced WC-2 binding at the *frq* promoter and loss of robust rhythmic *frq* transcription (Figure 8).

**Figure 8.**
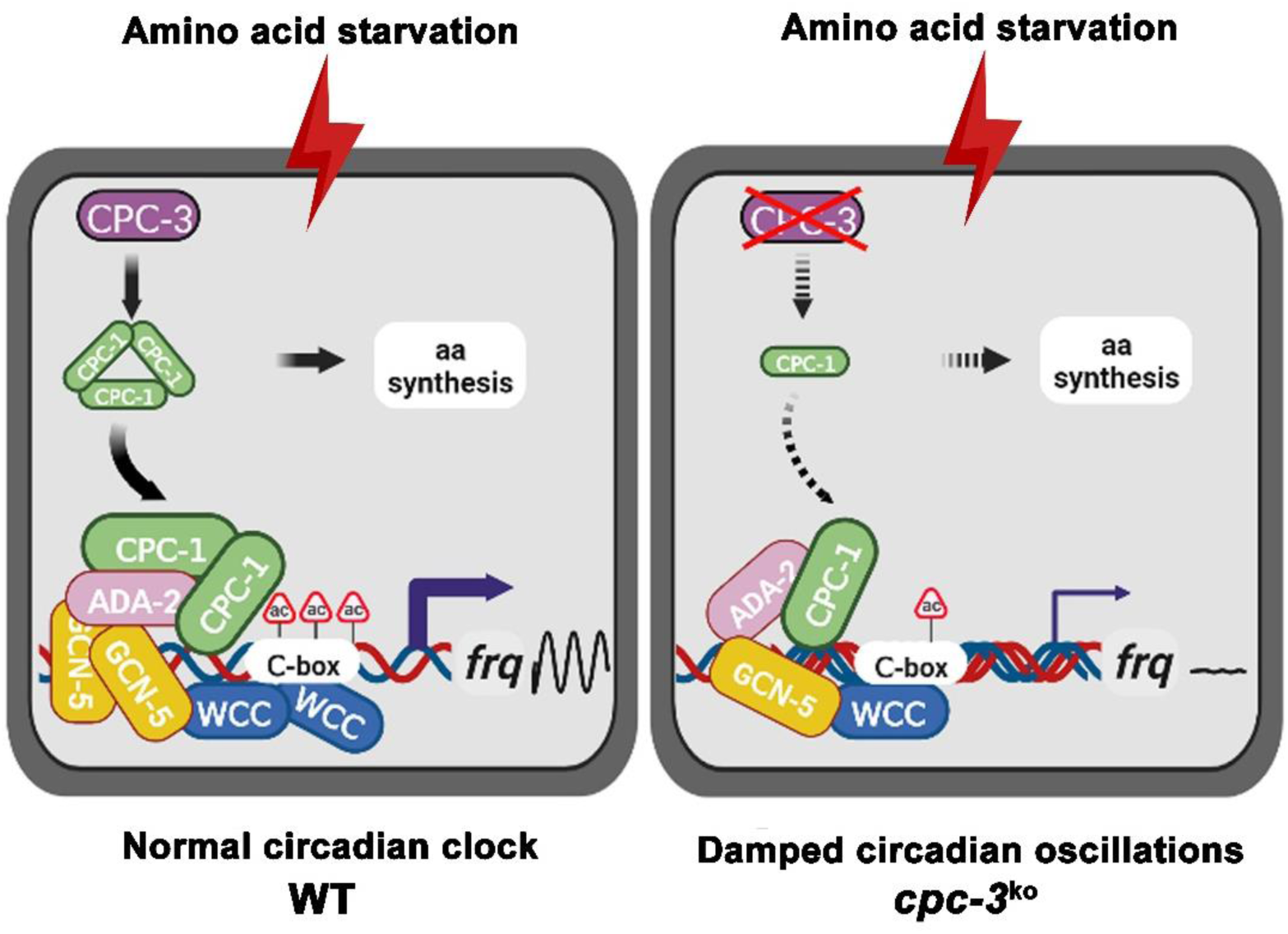
Model showing the role of CPC-3 and CPC-1 in maintaining the *Neurospora* circadian rhythm in response to amino acid starvation. CPC-3 and CPC-1 signaling pathway was activated by amino acid starvation and would recruit the histone acetyltransferase GCN-5 containing SAGA complex to promote the histone acetylation levels, which permitted rhythmic WC-2 binding at the *frq* promoter (Left). Disruption of the CPC-3 and CPC-1 signaling pathway resulted in decreased histone acetylation levels of the *frq* gene promoter, reduced WC-2 binding and damped circadian oscillations in response to the amino acid starvation stress (right).

Maintaining robust rhythmic gene expression and circadian activities in response to various environmental and nutrient stress is important for the health or survival of different organisms. Circadian clock synchronizes metabolic processes and systemic metabolite levels, while nutrients and energy signals also feedback to circadian clocks to adapt their metabolic state (Bass, 2012; Hurley et al., 2014; Klemz et al., 2017; Reinke & Asher, 2019). Amino acid starvation is known to inhibit the global translation efficiency through activating GCN2 mediated eIF2α phosphorylation, which conserves energy and allows cells to reprogram gene expression to relieve stress damage. It was recently shown that circadian clock control of GCN2 mediated eIF2α phosphorylation is necessary for rhythmic translation initiation in *Neurospora* (Ding et al., 2021; Karki et al., 2020). However, it was previously unknown whether circadian clock would be affected by amino acid starvation stress. After activation of the GCN2 mediated eIF2α phosphorylation by amino acid starvation, a subset of transcripts containing overlapping upstream open reading frames (uORFs) in their 5’ untranslated region (5’ UTR) are efficiently translated, including the yeast transcription factor GCN4, the *Neurospora* CPC-1 and their mammalian ortholog ATF4, which activate the transcription of various amino acid biosynthetic genes (Hinnebusch, 1984; Paluh, Orbach, Legerton, & Yanofsky, 1988; Vattem & Wek, 2004). Our ChIP results showed that CPC-1 could bind to the region close to C-box at *frq* promoter to activate *frq* transcription (Figure 4A). WCC binding at the *frq* C-box region slightly decreased under normal condition (Figure 2–figure supplement 1A), and dramatically decreased under amino acid starvation in the *cpc-1^KO^* strain (Figure 2D), suggesting that CPC-1 cooperates with WCC to promote *frq* transcription in response to amino acid starvation.

Amino acid starvation was able to affect gene expression by regulating chromatin modifications. Mammalian transcription factor ATF2 was reported to promote the modification of the chromatin structure in response to amino acid starvation to enhance the transcription of numbers of amino acid-regulated genes (Bruhat et al., 2007). Amino acid starvation was also shown to induce reactivation of silenced transgenes and latent HIV-1 provirus by down-regulation of histone deacetylase 4 in mammalian cells (Palmisano et al., 2012). Here we found that amino acid starvation suppressed clock gene *frq* expression by decreasing the histone acetylation levels, and activated GCN2 signaling pathway results in recruitment of the histone acetyltransferase SAGA complex to the *frq* promoter through its interaction with CPC-1 to re-establish a proper histone acetylation state to relieve the repression (Figures 3 and 4). Although histone acetylation and deacetylation rhythms have been reported in mammalian cells (Papazyan et al., 2016; Takahashi, 2017), their function is unclear in *Neurospora*. Here we demonstrated the important role of GCN-5 in regulating the rhythmic histone acetylation and WC-2 binding rhythms at the *frq* promoter (Figure 5). Our study unveiled an unsuspected link between nutrient limitation and circadian clock function mediated by the GCN2 signaling pathway. These results provide key insights into the epigenetic regulatory mechanisms of circadian gene expression during amino acid starvation.

The nutrient-sensing GCN2 signaling pathway is conserved in eukaryotic cells from yeast to mammals. In the budding yeast *Saccharomyces cerevisiae* and the filamentous fungus *Neurospora crassa*, the GCN2 kinase responds to nutrient deprivation, whereas it phosphorylates eIF2α and upregulates the master transcription factors GCN4 and CPC-1, respectively, that activate amino acid biosynthetic genes (Ebbole et al., 1991; Hinnebusch, 2005). In mammalian cells, several GCN2-related kinases can phosphorylate eIF2α in response to various stress conditions, triggering the integrated stress response (ISR) (Costa-Mattioli & Walter, 2020; Donnelly, Gorman, Gupta, & Samali, 2013). Interestingly, different from the normal circadian period of *cpc-3^KO^* strain in *Neurospora* without amino acid starvation (Figures 1A, Figure 1–figure supplement 1 and Figure 1–figure supplement 3), it was reported that GCN2 modulated circadian period by phosphorylation of eIF2α in mammals under normal condition, but it was unknown whether GCN2 was involved in circadian regulation of metabolism under nutrient limitation (Pathak et al., 2019). Our results suggest that the GCN2 signaling pathway is required for maintaining circadian clock under amino acid starvation, which is important for robust rhythmic expression of metabolic genes (Figure 7). Time-restricted feeding prevents obesity and metabolic syndrome through circadian related mechanisms(Chaix, Manoogian, Melkani, & Panda, 2019), but how eating pattern affects circadian regulation is under-studied. Based on the conservation of the GCN2 signaling pathway, our results indicated that GCN2 may play an important role in mediating circadian regulation of metabolism during nutrient limitation caused by time-restricted feeding in mammals. Together, our studies suggest a conserved role of GCN2 signaling pathway in maintaining the robustness of circadian clock under nutrient starvation in eukaryotes.

## Materials and Methods

### Strains, culture conditions, and race tube assay

The 87-3 (*bd*, a) and 301-6 (*bd, his-3^-^,* A) strain was used as the wild-type strain in this study. *cpc-3^KO^* (NCU01187), *cpc-1^KO^* (NCU04050), *gcn-5^KO^* (NCU10847), *ada-2^KO^* (NCU04459) and *hda-1^KO^* (NCU01525) strains were obtained from the Fungal Genetic Stock Center and were crossed with a *bd* strain to create the *bd;cpc-3^KO^*, *bd;cpc-1^KO^*, *bd;gcn-5^KO^*, *bd;ada-2^KO^* and *bd;hda-1^KO^* strains (Colot et al., 2006). Constructs with the *cpc-1* promoter driving expression of Myc.CPC-1 were introduced into the *cpc-1^KO^* strains at the *his-3* (NCU03139) locus by homologous recombination. Constructs with the *cfp* promoter driving expression of Flag.ADA-2 were introduced into the 301-6, cfp-Myc.CPC-1 or 301-6, cfp-Myc.GCN-5 strains by random insertion with nourseothricin selection(L. He et al., 2020). Positive transformants were identified by western blot analyses, and homokaryon strains were isolated by microconidia purification using 5 µm filters.

Liquid cultures were grown in minimal media (1x Vogel’s, 2% glucose). For rhythmic experiments, *Neurospora* was cultured in petri dishes in liquid medium for 2 days. The *Neurospora* mats were cut into discs and transferred into medium-containing flasks and were harvested at the indicated time points.

The medium for race tube assay contained 1x Vogel’s salts, 0.1% glucose, 0.17% arginine, 50 ng/mL biotin, and 1.5% agar. After entrainment of 24 hours in the constant light condition, race tubes were transferred to constant darkness conditions and marked every 24 hours. The circadian period of the *Neurospora* strain could be calculated according to the ratio between the distance of marked conidia band positions and the distance of conidiation bands.

### Luciferase reporter assay

The luciferase reporter assay was performed as reported previously (Gooch et al., 2008; Larrondo et al., 2015; X. Liu et al., 2017). The luciferase reporter construct (*frq*-*luc*) containing the luciferase gene under the control of the *frq* promoter, was introduced into 301-6, *cpc-3^KO^* and *cpc-1^KO^* strains by transformation. The luciferase reporter construct (FRQ-LUC) containing a luciferase fused to the C terminus of the FRQ protein, was introduced into *gcn-5^KO^* and *ada-2^KO^* strains by crossing. Firefly luciferin (final concentration of 50 μM) was added to autoclaved FGS-Vogel’s medium containing 1x FGS (0.05% fructose, 0.05% glucose, 2% sorbose), 1x Vogel’s medium, 50 μg/L biotin, and 1.8% agar. Conidia suspension was placed on autoclaved FGS-Vogel’s medium and grown in constant light overnight. The cultures were then transferred to constant darkness, and luminescence was recorded in real time using a LumiCycle after 1 day in DD. The data were then normalized with LumiCycle Analysis software by subtracting the baseline luciferase signal, which increases as cell grows.

### Protein analysis

Protein extraction, quantification, and western blot analyses were performed as previously described (X. Liu et al., 2017). Briefly, tissue was ground in liquid nitrogen with a mortar and pestle and suspended in ice-cold extraction buffer (50 mM HEPES (pH 7.4), 137 mM NaCl, 10% glycerol) with protease inhibitors (1 μg/mL Pepstatin A, 1 μg/mL Leupeptin, and 1 mM PMSF). After centrifugation, protein concentration was measured using protein assay dye (Bio-Rad). For western blot analyses, equal amounts of total protein (40 μg) were loaded in each protein lane of 7.5% or 10% SDS-PAGE gels containing a ratio of 37.5:1 acrylamide/bisacrylamide. After electrophoresis, proteins were transferred onto PVDF membranes, and western blot analyses were performed. Western blot signals were detected by X-ray films and scanned for quantification.

To detect the phosphorylation levels of WC-1 and WC-2, PPase inhibitors (25mM NaF, 10mM Na_4_P_2_O_7_.10H_2_O, 2mM Na_3_VO_4_, 1mM EDTA) were made fresh and added to the protein extraction buffer. Proteins were loaded in each protein lane of 7.5% SDS-PAGE gels containing a ratio of 149:1 acrylamide/bisacrylamide.

### RNA analysis

RNA was extracted with Trizol and further purified with 2.5 M LiCl as described previously(X. Liu et al., 2017). For northern blot analysis, equal amounts of total RNA (20 μg) were loaded onto agarose gels. After electrophoresis, the RNA was transferred onto nitrocellulose membrane. The membrane was probed with [^32^P] UTP (PerkinElmer)-labelled RNA probes specific for *frq*. RNA probes were transcribed in vitro from PCR products by T7 RNA polymerase (Invitrogen, AM1314M) with the manufacturer’s protocol. The *frq* primers used for the template amplification were shown in Figure 7–Table supplement 11.

For RT-qPCR, the cultures of WT and *cpc-1^KO^* strains were collected at the indicated time points in constant darkness in liquid growth medium (1x Vogel’s, 2% glucose). RT-qPCR were performed as previously described(Cui et al., 2020). Each RNA sample (1 μg) was subjected to reverse transcription with HiScript II reverse transcriptase (Vazyme, R223), and then amplified by real-time PCR (Bio-Rad, CFX96). For RT-qPCR, primers target the coding genes of *frq*(NCU02265), *his-3* (NCU03139), *ser-2* (NCU01439) and *trp-3* (NCU08409) were designed, and the β-tubulin was used as an internal control. The primers used for RT-qPCR were shown in Figure 7–Table supplement 1.

### Generation of antiserum against CPC-1

Two CPC-1 peptides (SELDLLDFATFDGG and RDKPLPPIIVEDPS) were synthesized and used as the antigens to generate rabbit polyclonal antisera (ABclonal) as described previously (Cui et al., 2020; Zhou et al., 2013).

### Immunoprecipitation (IP) analysis

Immunoprecipitation analyses were performed as previously described (Cao et al., 2018; Cheng, Yang, Heintzen, & Liu, 2001). Briefly, *Neurospora* proteins were extracted as described above. For each immunoprecipitation reaction, 2 mg protein and 2 μL c-Myc (TransGen, HT101), 2 μL Flag (Sigma, F1804) or 1 μL WC-2 antibody (Cheng, Yang, Heintzen, et al., 2001) were used. After incubation with antibody for 3 hours, 40 μL GammaBind G Sepharose beads (GE Healthcare, 17061801) were added, and samples were incubated for 1 hour. Immunoprecipitated proteins were washed three times using extraction buffer before western blot analysis. IP experiments were performed using cultures harvested in constant light.

### Chromatin immunoprecipitation (ChIP) analysis

ChIP assays were performed as previously described (Cui et al., 2020; Zhou et al., 2013) with 2 mg protein used for each immunoprecipitation reaction. The chromatin immunoprecipitation reaction was carried out with 2 μL WC-2 (Cheng, Yang, Heintzen, et al., 2001), H2B (Abcam,ab1790) or H3ac (Millipore, 06-599) antibody. Immunoprecipitated DNA was quantified by real-time qPCR. Occupancies were normalized by the ratio of ChIP to Input DNA. ChIP was performed using 5 μL c-Myc monoclonal antibody (TransGen, HT101) to examine occupancy of Myc.CPC-1 or Myc.GCN-5. Occupancies of ChIP were normalized using IgG. The primers used for ChIP-qPCR were shown in Figure 7–Table supplement 1.

### RNA-seq analysis

The WT and *cpc-1^KO^* strains were cultured with or without 12 mM 3-AT treatment in constant light. Total RNAs were extracted using Trizol reagents. Libraries were prepared according to the manufacturer’s instructions and analyzed using 150bp paired-end Illumina sequencing (Annoroad Gene Technology, Beijing). After sequencing, the raw data was treated and mapped to the genome of *Neurospora crassa* and transformed into expression value. The gene expression levels were scored by fragments per kb per million (FPKM) method. The differences in gene expression between samples was compared by comparing FPKM values, and those with fold change more than 1 (FDR<0.1) were thought to be differentially expressed genes (DEGs). The functional category enrichments, including Gene Ontology (GO) and KEGG terms, were analyzed. The KEGG pathway enrichment was evaluated based on hypergeometric distribution, and the R package ‘ggplot2’ version 3.3.6 was used to visualize the enrichment results.

### Quantification and statistical analysis

Quantification of western blot data were performed using Image J software. Error bars are standard deviations for chromatin immunoprecipitation assays from at least three independent technical experiments, and standard error of means for race tube assays from at least three independent biological experiments, unless otherwise indicated. Statistical significance was determined by Student’s t-test.

## Acknowledgements

We thank Dr. Linqi Wang, Dr. Wenbing Yin and members of our lab for assistance and discussions. We thank Dr. Luis Larrondo for providing the translational fused FRQ-LUC plasmid and strain. This work was supported by grants from National Natural Science Foundation of China (32170092, 31970079), National Key Research and Development Program of China (2021YFA0911300), Strategic Priority Research Program of the Chinese Academy of Sciences (XDA28030402), Beijing Natural Science Foundation (5202020), and CAS Interdisciplinary Innovation Team to Xiao Liu; National Natural Science Foundation of China (32200056) to Xiaolan Liu; National Key Research and Development Program of China (2018YFA0900500) and National Natural Science Foundation of China (32170560) to Qun He; National Institutes of Health (R35 GM118118) and the Welch Foundation (I-1560) to Yi Liu. The funders had no role in study design, data collection and analysis, decision to publish, or preparation of the manuscript.

## Author Contributions

Conceptualization, Xiao Liu, Xiaolan Liu, and Yulin Yang; Investigation, Xiaolan Liu, Yulin Yang, Yue Hu, Jingjing Wu, Xihui Gan, Chuqiao Han, Qiaojia Lu and Shaohua Qi; Data analysis, Xiao Liu, Xiaolan Liu, Yulin Yang, Yue Hu, Qiaojia Lu, Qun He, Jinhu Guo and Yi Liu; Validation, Xiaolan Liu, Yulin Yang and Yue Hu; Funding acquisition, Xiao Liu, Xiaolan Liu, Qun He and Yi Liu; Resources, Xiao Liu, Qun He, Jinhu Guo and Yi Liu; Writing, Xiao Liu, Yi Liu and Xiaolan Liu; Supervision, Xiao Liu.

## Data and materials availability

RNA Sequencing data have been deposited in GEO under accession code GSE220169. Materials are available from the corresponding authors upon reasonable request.

## Additional source data included

**Figure 1 - source data 1.** Scan of western blot probed for FRQ protein in Figure 1C.

**Figure 1 - source data 2.** Scan of Northern blot probed for *frq* mRNA in Figure 1D.

**Figure 2 - source data 1.** Scan of western blot probed for WC-1 and WC-2 proteins in WT and *cpc- 3^KO^* strains with 3-AT in Figure 2A.

**Figure 2 - source data 2.** Scan of western blot probed for WC-1 and WC-2 proteins in WT and *cpc- 1^KO^* strains with 3-AT in Figure 2B.

**Figure 4 - source data 1.** Scan of western blot probed for Flag.ADA-2 and Myc.GCN-5 proteins in Figure 4B.

**Figure 4 - source data 2.** Scan of western blot probed for Flag.ADA-2 and Myc.CPC-1 proteins in Figure 4C.

**Figure 5 - source data 1.** Scan of western blot probed for FRQ protein in Figure 5C.

**Figure 5 - source data 2.** Scan of Northern blot probed for *frq* mRNA in Figure 5D.

**Figure 1 - figure supplement 1 - source data 1.** Scan of western blot probed for phosphorylation of eIF2α protein in Figure 1 - figure supplement 1A.

**Figure 1 - figure supplement 1 - source data 1.** Scan of western blot probed for CPC-1 protein in Figure 1 - figure supplement 1B.

**Figure 1 - figure supplement 3 - source data 1.** Scan of western blot probed for FRQ protein in Figure 1 - figure supplement 3B.

**Figure 1 - figure supplement 3 - source data 2.** Scan of Northern blot probed for *frq* mRNA in Figure 1 - figure supplement 3C.

**Figure 2 - figure supplement 1 - source data 1.** Scan of western blot probed for phosphorylation of WC-1 and WC-2 proteins in WT and *cpc-3^KO^* strains with 3-AT in Figure 2 - figure supplement 1B.

**Figure 2 - figure supplement 1 - source data 2.** Scan of western blot probed for phosphorylation of WC-1 and WC-2 proteins in WT and *cpc-1^KO^* strains with 3-AT in Figure 2 - figure supplement 1C.

**Figure 4 - figure supplement 1 - source data 1.** Scan of western blot probed for Myc.CPC-1, WC-1 and WC-2 proteins in Figure 4 - figure supplement 1A.

**Figure 4 - figure supplement 1 - source data 2.** Scan of western blot probed for Myc.GCN-5, WC-1 and WC-2 proteins in Figure 4 - figure supplement 1B.

**Figure 5 - figure supplement 1 - source data 1.** Scan of western blot probed for FRQ protein in Figure 5 - figure supplement 1C.

---

**Figure 1–figure supplement 1.**
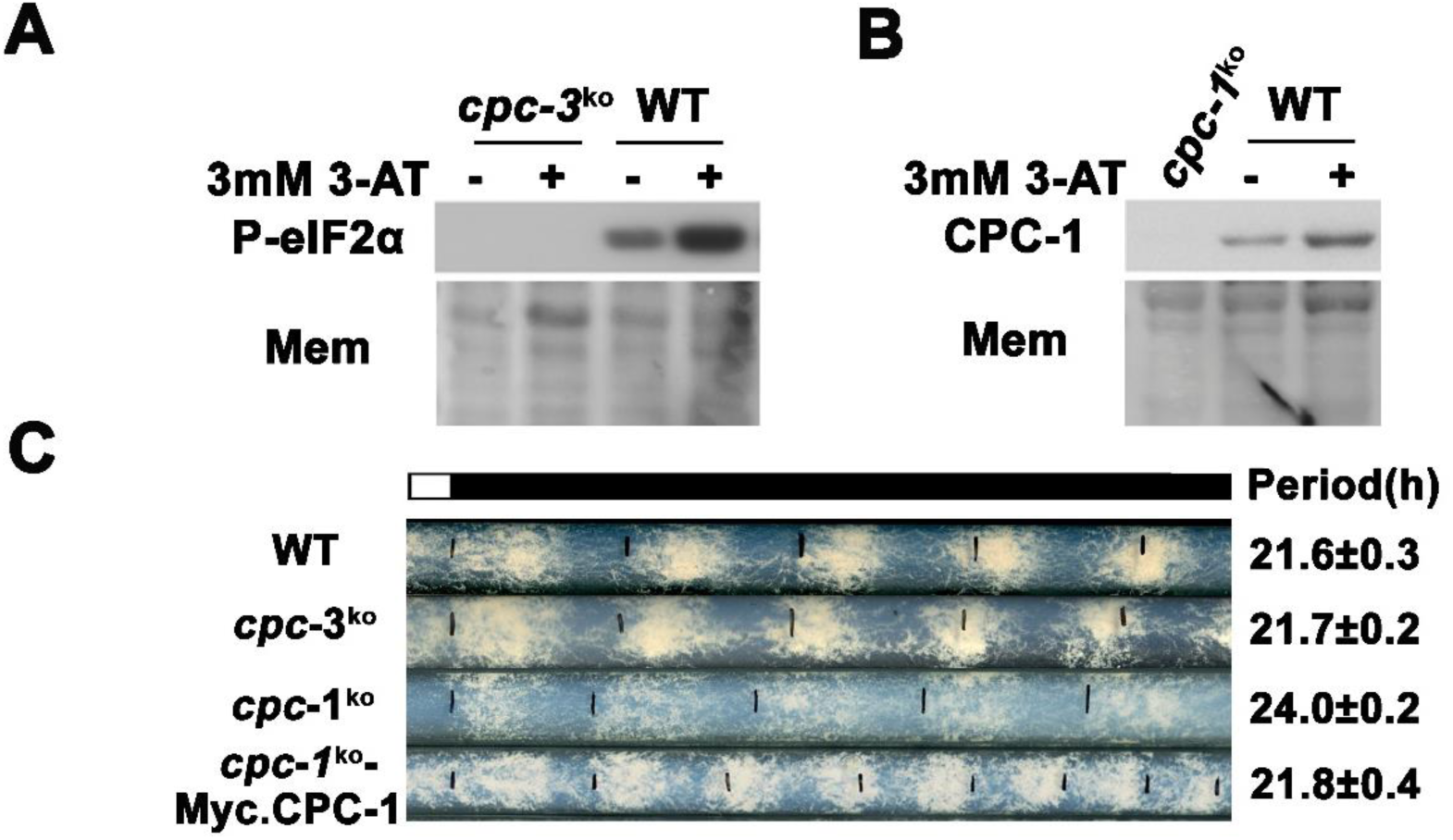
Circadian rhythm of *cpc-3^KO^* and *cpc-1^KO^* strain. (A) eIF2α was phosphorylated by CPC-3 in the presence of 3 mM 3-AT. Western blot of proteins from WT and *cpc-3^KO^* strains treated (+) or not (−) with 3-AT using P-eIF2α antibody. Mem: stained membrane (loading control). (B) CPC-1 was activated in the presence of 3 mM 3-AT. Western blot of proteins from WT and *cpc-1^KO^* strains harvested in the indicated hours after 3-AT treatment using CPC-1 antibody. Mem: stained membrane. (C) Race tube assay showing that expression of Myc-CPC-1 rescued the prolonged circadian period of *cpc-1^KO^* strain.

**Figure 1–figure supplement 2.**
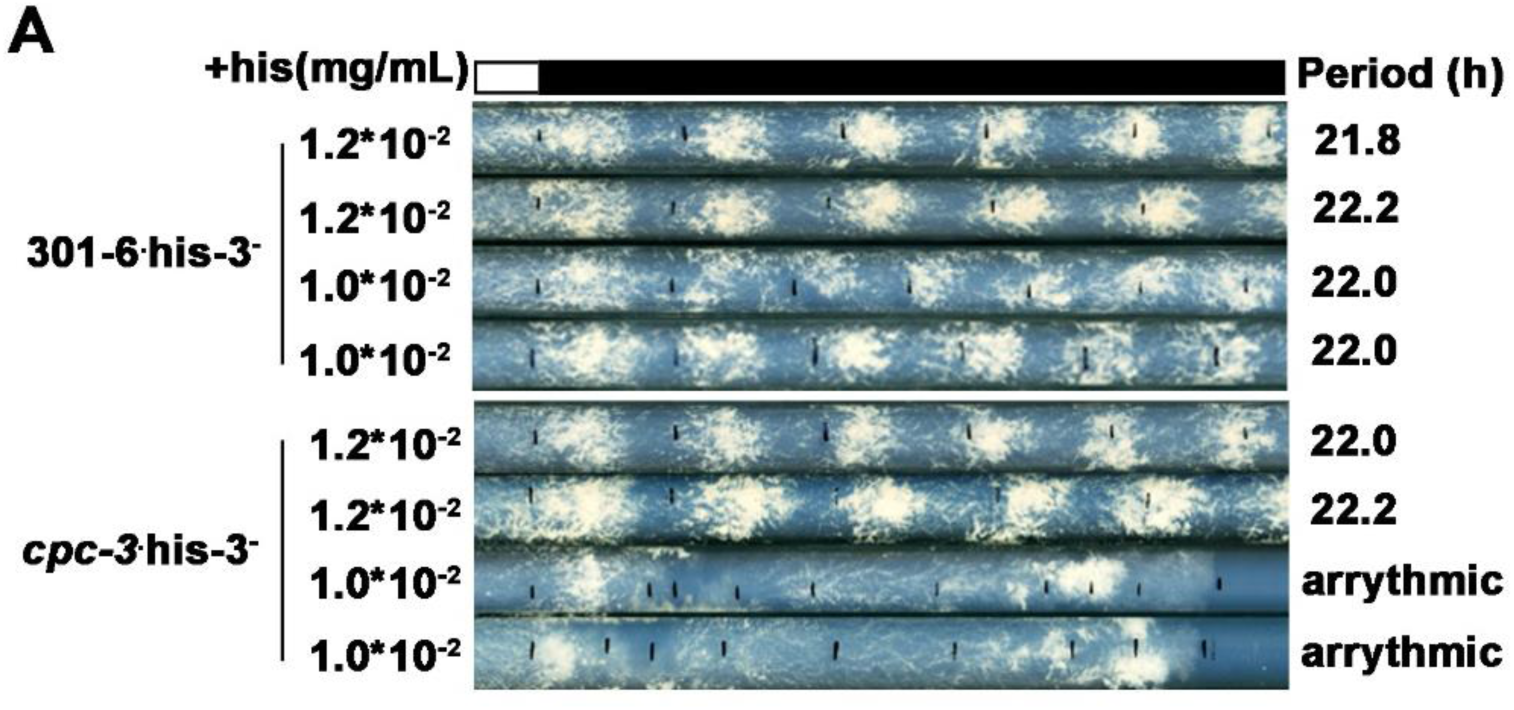
CPC-3 is required for robust circadian conidiation rhythm under histidine starvation. (A) Race tube assay showed that the *his-3^-^* strain could grow with 1.0*10^-2^ mg/mL histidine, and exhibited normal circadian conidiation rhythm, however, the *cpc-3 ^KO^ his-3^-^* strains lost circadian conidiation rhythm with limited histidine at the concentration of 1.0*10^-2^ mg/mL.

**Figure 1–figure supplement 3.**
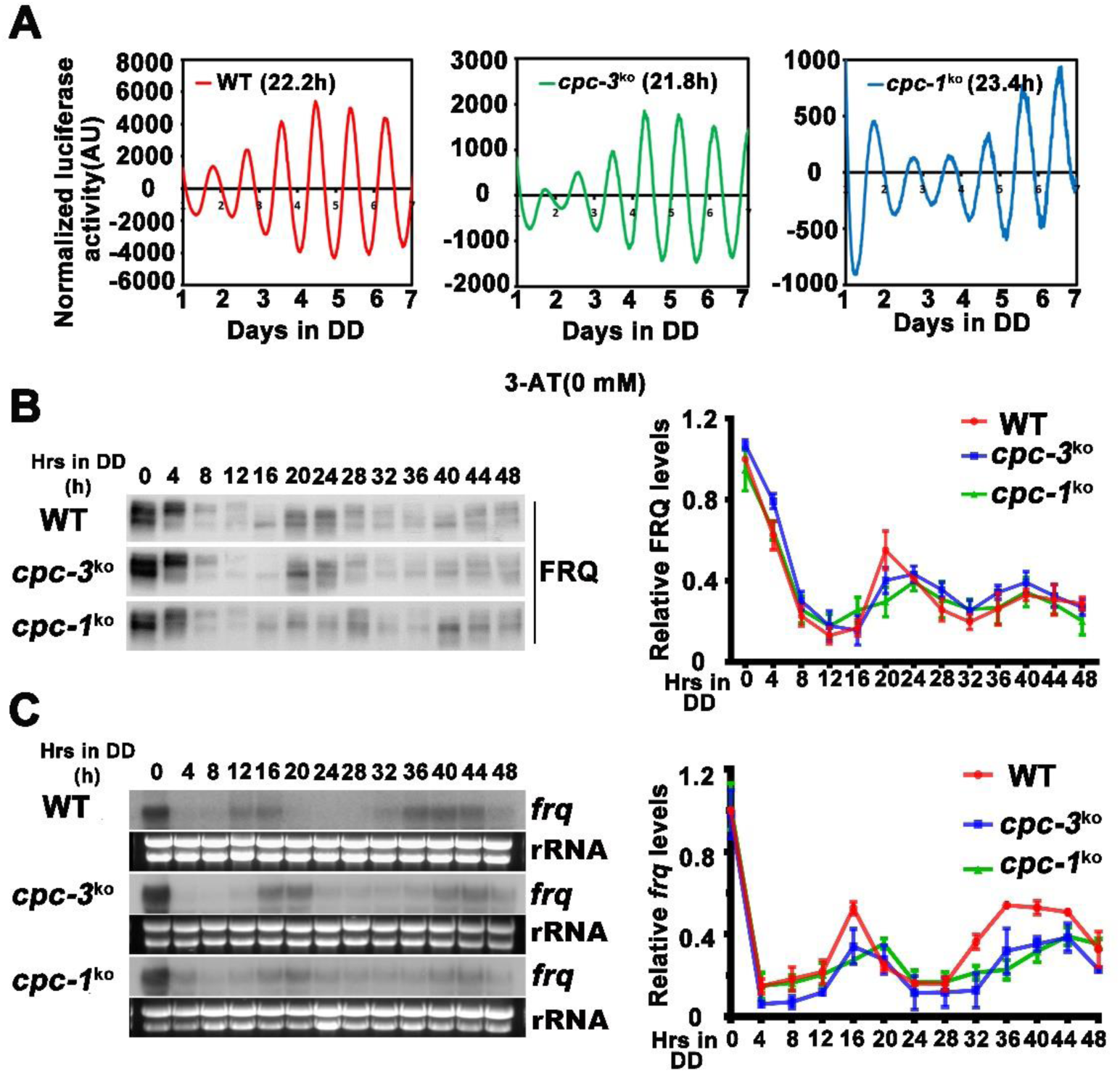
Rhythmic *frq* expression of *cpc-3^KO^* and *cpc-1^KO^* strain. (A) Luciferase reporter assay showing normal circadian period of *cpc-3^KO^* strain and prolonged circadian period of *cpc-1^KO^* strain. A *frq-luc* transcriptional fusion construct was expressed in *cpc-3^KO^* and *cpc-1^KO^* strains grown on the FGS-Vogel’s medium, and the luciferase signal was recorded using a LumiCycle in constant darkness (DD) for more than 7 days. Normalized data with the baseline luciferase signals subtracted are shown. (B) Western blot assay showing the rhythmic expression of FRQ protein in the *cpc-3^KO^* and *cpc-1^KO^* strains at the indicated time points in DD. The left panel showing that protein extracts were isolated from WT, *cpc-3^KO^* and *cpc-1^KO^* strains grown in a circadian time course in DD and probed with FRQ antibody. The right panel showing that the densitometric analyses of the results of three independent experiments. (C) Northern blot assay showing the rhythmic expression of *frq* mRNA in the *cpc-3^KO^* and c*pc-1^KO^* strains at the indicated time points in DD. The densitometric analyses of the results from three independent experiments are shown on the right panel.

**Figure 2–figure supplement 1.**
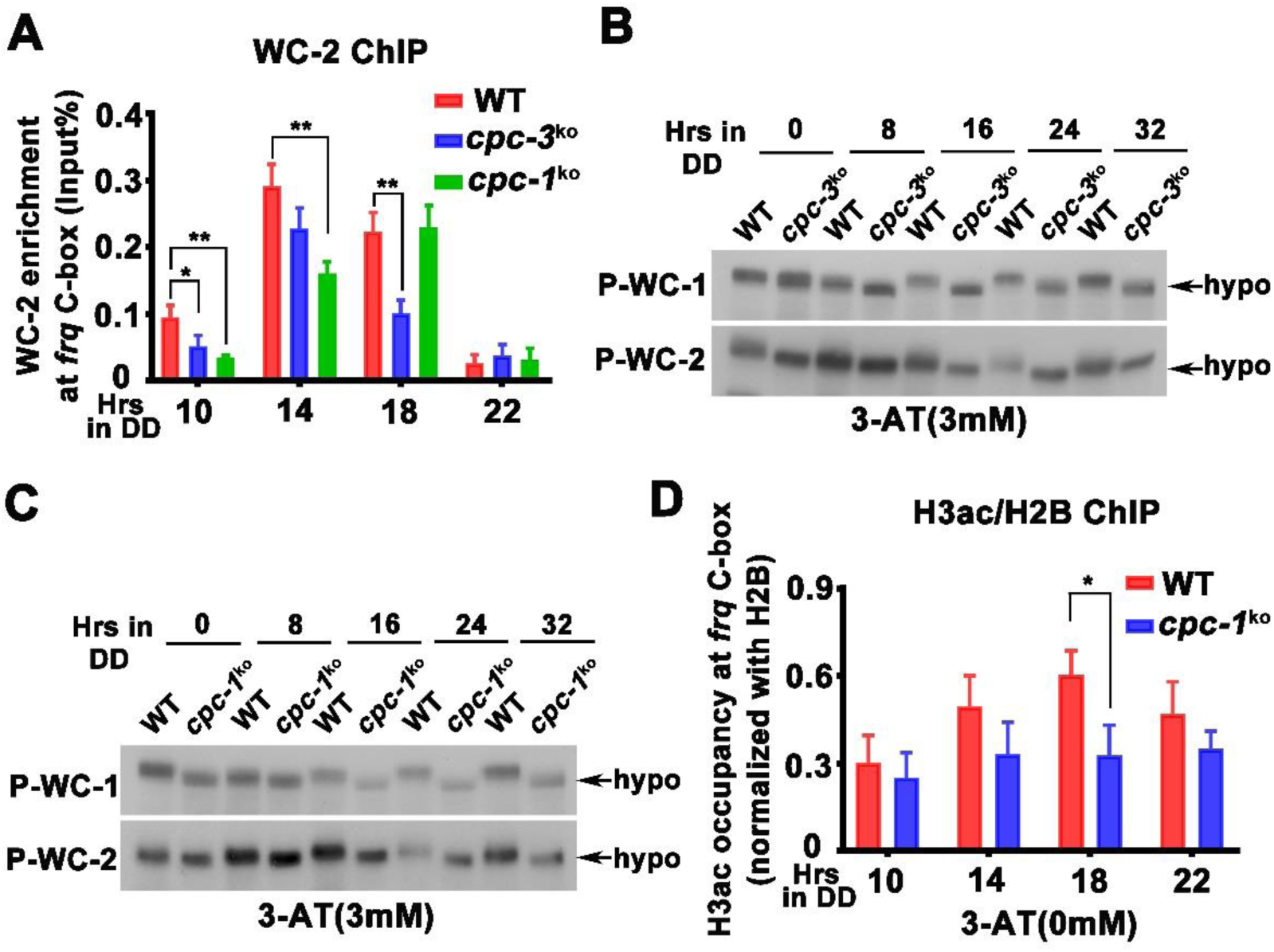
WCC binding and WCC phosphorylation levels at *frq* promoter in the *cpc-3^KO^* and *cpc-1^KO^* strain. (A) ChIP assay showing that deletion of *cpc-3* or *cpc-1* did not dramatically affect WC-2 binding at the promoter of *frq* gene without 3-AT. Samples were grown for the indicated number of hours in DD prior to harvesting and processing for ChIP using WC-2 antibody. Occupancies were normalized by the ratio of ChIP to Input DNA. Error bars indicate standard deviation (n = 3). *p < 0.05; **p < 0.01; Student’s t test was used. (B-C) Western blot assay showing that WCC was hypo-phosphorylated in the *cpc-3^KO^* (B) and *cpc-1^KO^* (C) strains in the presence of 3 mM 3-AT. Proteins were extracted by protein extraction buffer with PPase inhibitors and loaded in each protein lane of 7.5% SDS-PAGE gels containing a ratio of 149:1 acrylamide/bisacrylamide. The faster mobility shift indicated hypo-phosphorylation levels of WC-1 and WC-2. (D) ChIP assay showing slightly decreased histone H3ac levels at the promoter of *frq* gene in the *cpc-1^KO^* strain without 3-AT at the indicated time points in DD. Error bars indicate standard deviations (n=3). *P<0.05; Student’s t test was used.

**Figure 4–figure supplement 1.**
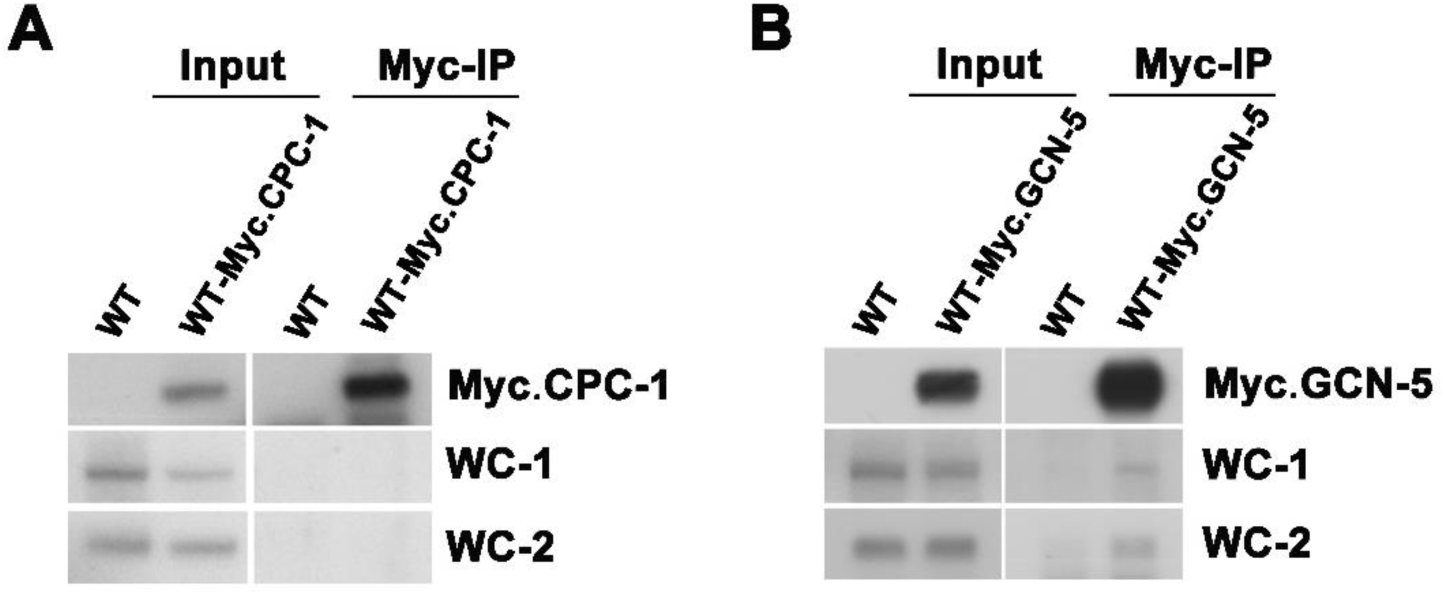
GCN-5 interacts with WCC. (A) IP assay showing that CPC-1 did not interact with WCC. Myc-CPC-1 was expressed in the WT strain and immunoprecipitation was performed using Myc antibody. (B) IP assay showing that GCN-5 interacted with WC-1 and WC-2. Myc-GCN-5 was expressed in the WT strain and immunoprecipitation was performed using Myc antibody.

**Figure 5–figure supplement 1.**
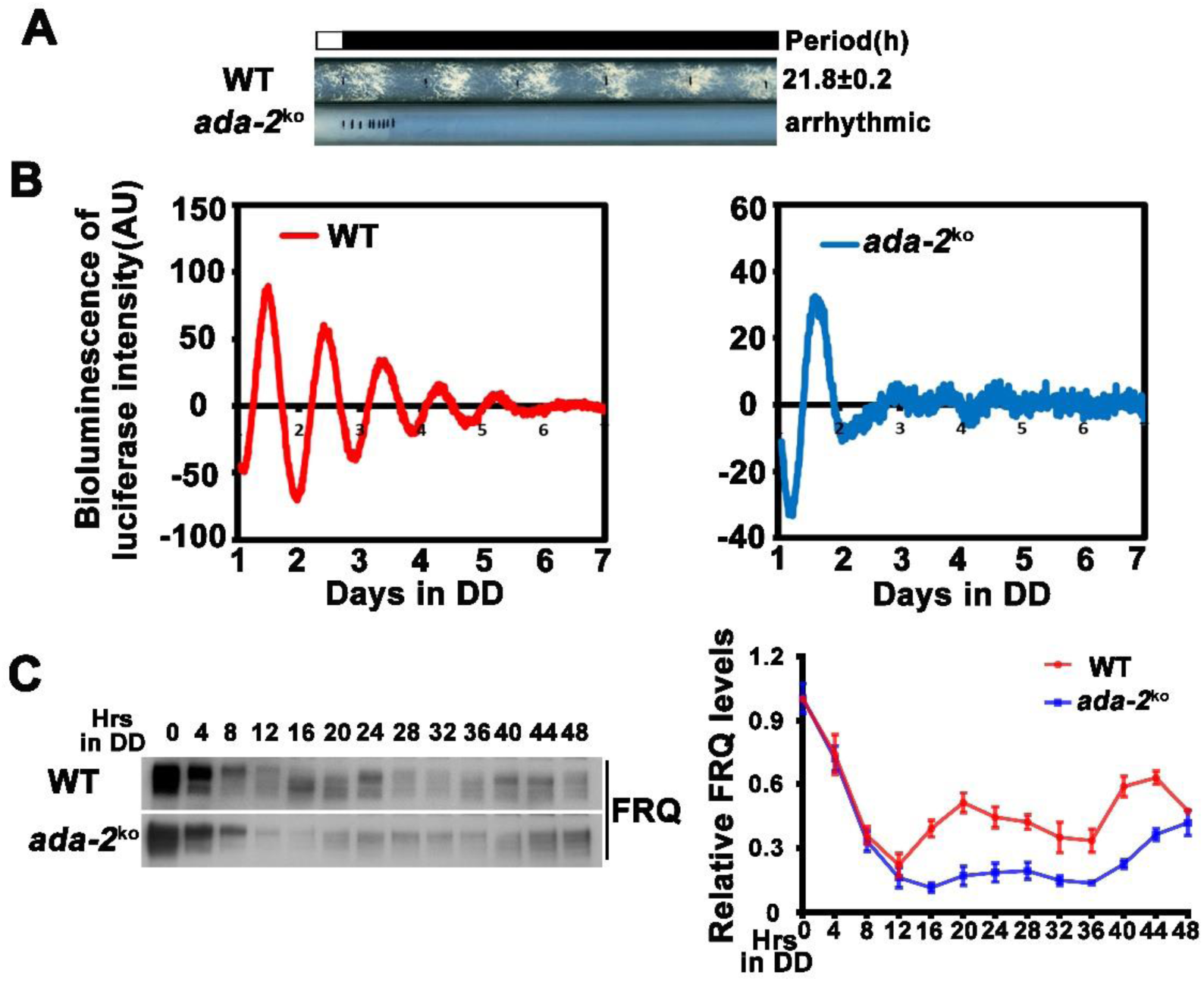
ADA-2 is required for circadian rhythm by regulating rhythmic *frq* expression. (A) Race tube assay showing that the conidiation rhythm in *ada-2^KO^* strains was lost compared with wild-type strains. (B) Luciferase assay showing that the luciferase activity rhythm was impaired in the *ada-2^KO^* strain after one day transition from light to dark. A FRQ-LUC translational fusion construct was expressed in WT and *ada-2^KO^* strains, and the luciferase signal was recorded in DD for more than 7 days. Normalized data with the baseline luciferase signals subtracted are shown. (C) Western blot assay showing that rhythmic expression of FRQ protein was dampened in the *ada-2^KO^* strain. The right panel showing that the densitometric analyses of the results of three independent experiments.

**Figure 7–figure supplement 1.**
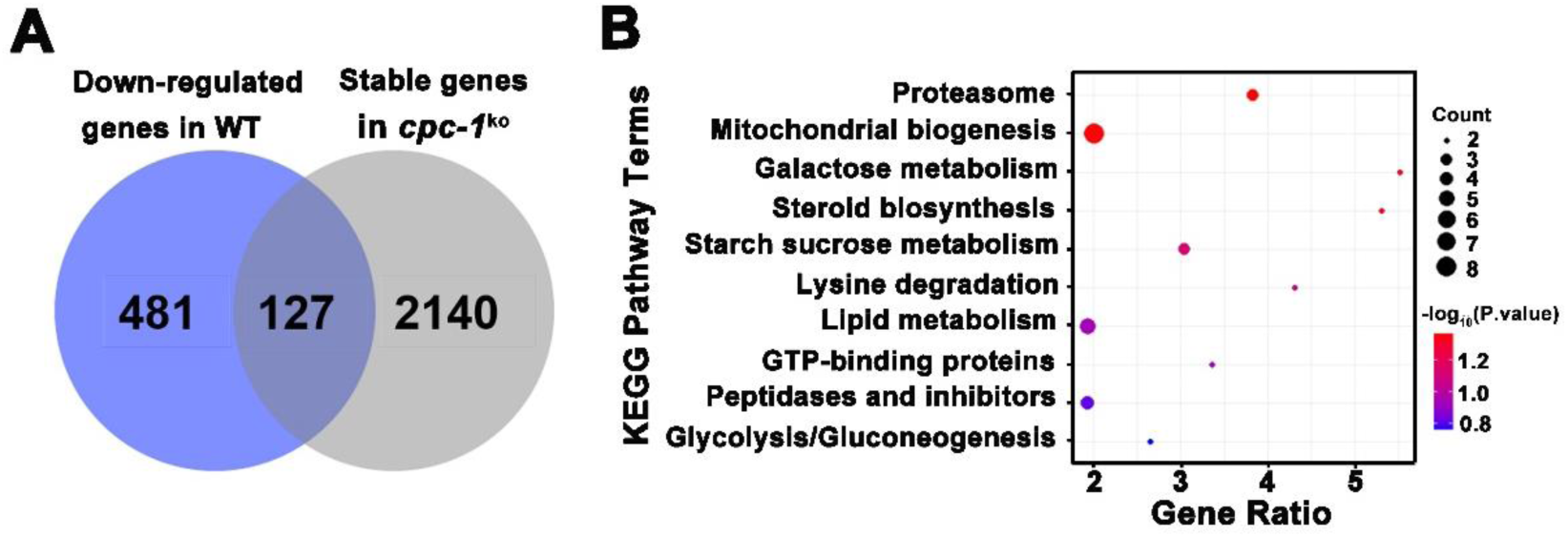
CPC-1 regulated metabolic genes under amino acid starvation. (A) Pie charts showing the overlaps of down-regulated genes in the WT strain, but unchanged genes in the *cpc-1^KO^* strains. (B) Gene functional enrichment analysis based on the mRNA level changes for the overlaps of down-regulated genes in the WT strain, but stable genes in the *cpc-1^KO^* strains.

**Figure 7–Table supplement 1.**
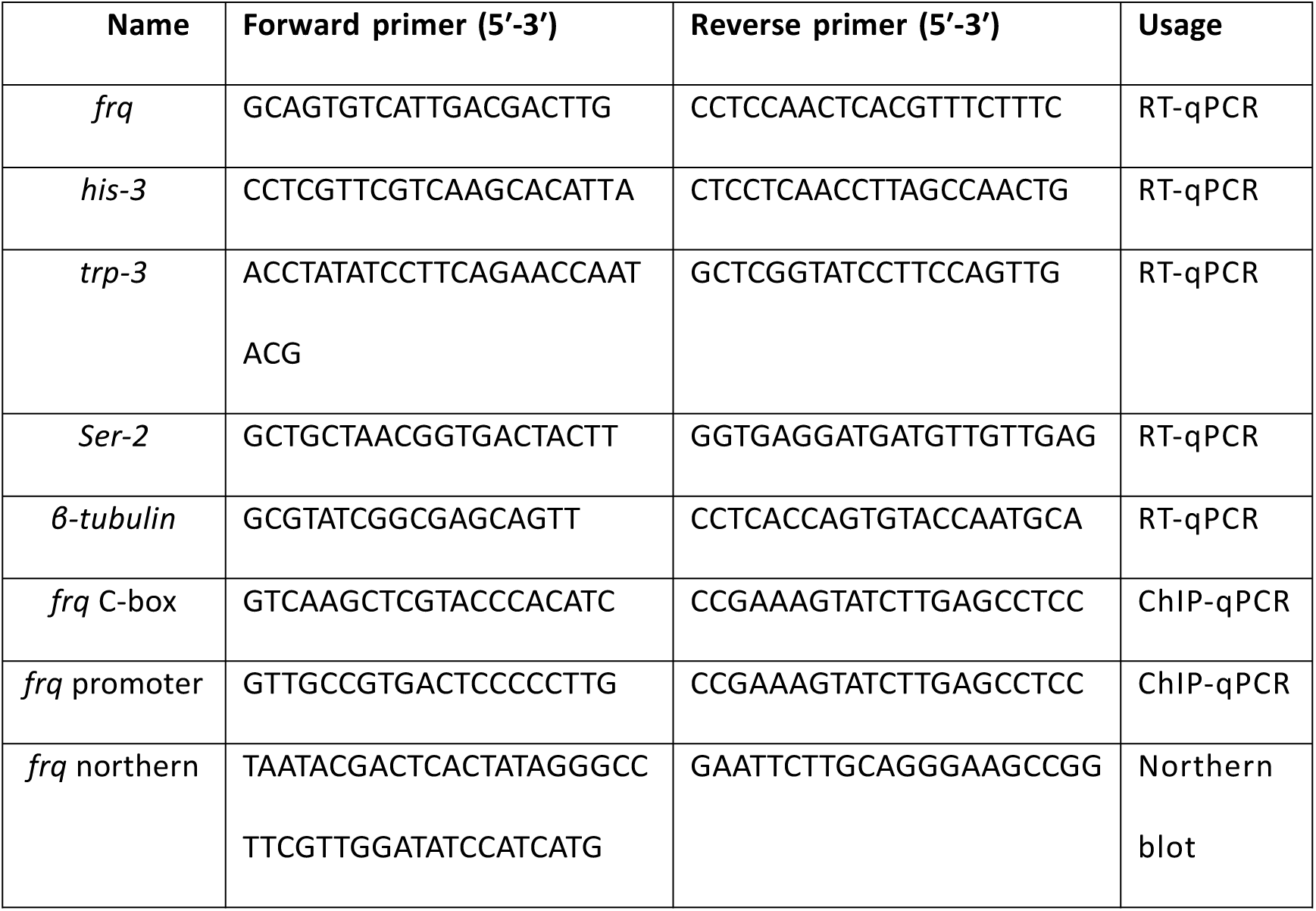
Primers used for ChIP-qPCR, Northern blot and RT-qPCR.

